# Plant photoreceptors and their signaling components compete for binding to the ubiquitin ligase COP1 using their VP-peptide motifs

**DOI:** 10.1101/568618

**Authors:** Kelvin Lau, Roman Podolec, Richard Chappuis, Roman Ulm, Michael Hothorn

## Abstract

Plants sense different parts of the sun’s light spectrum using specialized photoreceptors, many of which signal through the E3 ubiquitin ligase COP1. Photoreceptor binding modulates COP1’s ubiquitin ligase activity towards transcription factors. Here we analyze why many COP1-interacting transcription factors and photoreceptors harbor sequence-divergent Val-Pro (VP) peptide motifs. We demonstrate that VP motifs enable different light signaling components to bind to the WD40 domain of COP1 with various binding affinities. Crystal structures of the VP motifs of the UV-B photoreceptor UVR8 and the transcription factor HY5 in complex with COP1, quantitative binding assays and reverse genetic experiments together suggest that UVR8 and HY5 compete for the COP1 WD40 domain. Photoactivation of UVR8 leads to high-affinity cooperative binding of its VP domain and its photosensing core to COP1, interfering with the binding of COP1 to its substrate HY5. Functional UVR8 – VP motif chimeras suggest that UV-B signaling specificity resides in the UVR8 photoreceptor core, not its VP motif. Crystal structures of different COP1 – VP peptide complexes highlight sequence fingerprints required for COP1 targeting. The functionally distinct blue light receptors CRY1 and CRY2 also compete with downstream transcription factors for COP1 binding using similar VP-peptide motifs. Together, our work reveals that photoreceptors and their components compete for COP1 using a conserved displacement mechanism to control different light signaling cascades in plants.

## INTRODUCTION

Flowering plants etiolate in darkness, manifested by the rapid elongation of the embryonic stem, the hypocotyl, and closed and underdeveloped embryonic leaves, the cotyledons. Under light and upon photoreceptor activation, seedlings de-etiolate and display a photomorphogenic phenotype, characterized by a short hypocotyl and open green cotyledons, enabling a photosynthetic lifestyle (Gommers and Monte, 2018). The *constitutively photomorphogenic 1* (*cop1*) mutant displays a light-grown phenotype in the dark, including a short hypocotyl, and open and expanded cotyledons. *COP1* is thus a crucial repressor of photomorphogenesis (Deng et al., 1991). COP1 contains an N-terminal zinc-finger, a central coiled-coil, and a C-terminal WD40 domain, which is essential for proper COP1 function (Deng et al., 1992; McNellis et al., 1994). Light-activated phytochrome, cryptochrome and UVR8 photoreceptors inhibit COP1’s activity (von Arnim and Deng, 1994; Hoecker, 2017; Podolec and Ulm, 2018). Although COP1 can act as a stand-alone E3 ubiquitin ligase *in vitro* (Seo et al., 2003; Saijo et al., 2003), it forms higher-order complexes *in vivo*, for example with SUPPRESSOR OF PHYA-105 (SPA) proteins (Hoecker and Quail, 2001; Zhu et al., 2008; Ordoñez-Herrera et al., 2015). COP1 can also act as a substrate adaptor in CULLIN4 – DAMAGED DNA BINDING PROTEIN 1 (CUL4-DDB1)-based heteromeric E3 ubiquitin ligase complexes (Chen et al., 2010). These different complexes may modulate COP1’s activity towards different substrates (Ren et al., 2019). COP1 regulates gene expression and plays a central role as a repressor of photomorphogenesis by directly modulating the stability of transcription factors that control the expression of light-regulated genes (Lau and Deng, 2012; Podolec and Ulm, 2018). For example, the bZIP transcription factor ELONGATED HYPOCOTYL 5 (HY5) acts antagonistically with COP1 (Ang et al., 1998). COP1 binding to HY5 leads to its subsequent degradation via the 26S proteasome in darkness, a process that is inhibited by light (Osterlund et al., 2000).

In addition to HY5, other COP1 targets have been identified including transcriptional regulators, such as the HY5 homolog HYH (Holm et al., 2002), CONSTANS (CO) and other members of the BBX protein family (Jang et al., 2008; Liu et al., 2008; Khanna et al., 2009; Xu et al., 2016; Lin et al., 2018; Ordoñez-Herrera et al., 2018), and others such as LONG HYPOCOTYL IN FAR-RED (HFR1) (Jang et al., 2005; Yang et al., 2005) and SHI-RELATED SEQUENCE5 (SRS5) (Yuan et al., 2018). It has been suggested that specific Val-Pro (VP)-peptide motifs with a core sequence V-P-E/D-Φ-G, where Φ designated a hydrophobic residue, are able to bind the COP1 WD40 domain (Holm et al., 2001; Uljon et al., 2016). Deletion of regions containing the VP-peptide motifs result in loss of interaction of COP1 substrates with the COP1 WD40 domain (Holm et al., 2001; Jang et al., 2005; Datta et al., 2006). Recently, it has been reported that the human COP1 WD40 domain directly binds VP-motifs, such as the one from the TRIB1 pseudokinase that acts as a scaffold facilitating ubiquitination of human COP1 substrates (Uljon et al., 2016; Durzynska et al., 2017; Newton et al., 2018; Kung and Jura, 2019).

Arabidopsis photoreceptors for UV-B radiation (UV RESISTANCE LOCUS 8, UVR8), for blue light (cryptochrome 1 and 2, CRY1/CRY2) and for red/far-red light (phytochromes A-E), are known to repress COP1 activity in a light-dependent fashion (Yang et al., 2000; Wang et al., 2001; Yu et al., 2007; Favory et al., 2009; Jang et al., 2010; Lian et al., 2011; Liu et al., 2011; Zuo et al., 2011; Viczián et al., 2012; Huang et al., 2013; Lu et al., 2015; Sheerin et al., 2015; Yang et al., 2018). UVR8 itself contains a conserved C-terminal VP-peptide motif that is critical for UV-B signaling (Cloix et al., 2012; Yin et al., 2015). Moreover, overexpression of the UVR8 C-terminal 44 amino acids results in a cop-like phenotype (Yin et al., 2015). A similar phenotype has been observed when overexpressing the COP1-interacting CRY1 and CRY2 C-terminal domains (CCT) (Yang et al., 2000, 2001). Indeed, CRY1 and CRY2 also contain potential VP-peptide motifs within their CCT domains, but their function in blue-light signaling has not been established (Lin and Shalitin, 2003; Müller and Bouly, 2015). The presence of VP-peptide motifs in different light signaling components suggests that COP1 may use a common targeting mechanism to interact with downstream transcription factors and upstream photoreceptors. Here we present structural, quantitative biochemical and genetic evidence for a VP-peptide-based competition mechanism, enabling COP1 to play a crucial role in different photoreceptor pathways in plants.

## RESULTS

### The COP1 WD40 domain binds VP motifs from UVR8 and HY5

The WD40 domains of human and Arabidopsis COP1 can directly sense VP-containing peptides (Uljon et al., 2016). Such a VP-peptide motif can be found in the UVR8 C-terminus that is not part of the UV-B-sensing β-propeller domain (Figure 1A) (Kliebenstein et al., 2002; Rizzini et al., 2011; Christie et al., 2012; Wu et al., 2012), but is essential for UV-B signaling (Cloix et al., 2012; Yin et al., 2015). HY5 (Oyama et al., 1997), which is a COP1 target acting downstream of UVR8 in the UV-B signaling pathway (Ulm et al., 2004; Brown et al., 2005; Oravecz et al., 2006; Binkert et al., 2014), also contains a VP-peptide motif (Figure 1A) (Holm et al., 2001). UV-B absorption leads to UVR8 monomerization, COP1 binding and subsequent stabilization of HY5 (Favory et al., 2009; Rizzini et al., 2011; Huang et al., 2013). Mutation of the HY5 VP pair to alanine (AA) stabilizes the HY5 protein (Holm et al., 2001).

**Figure 1:**
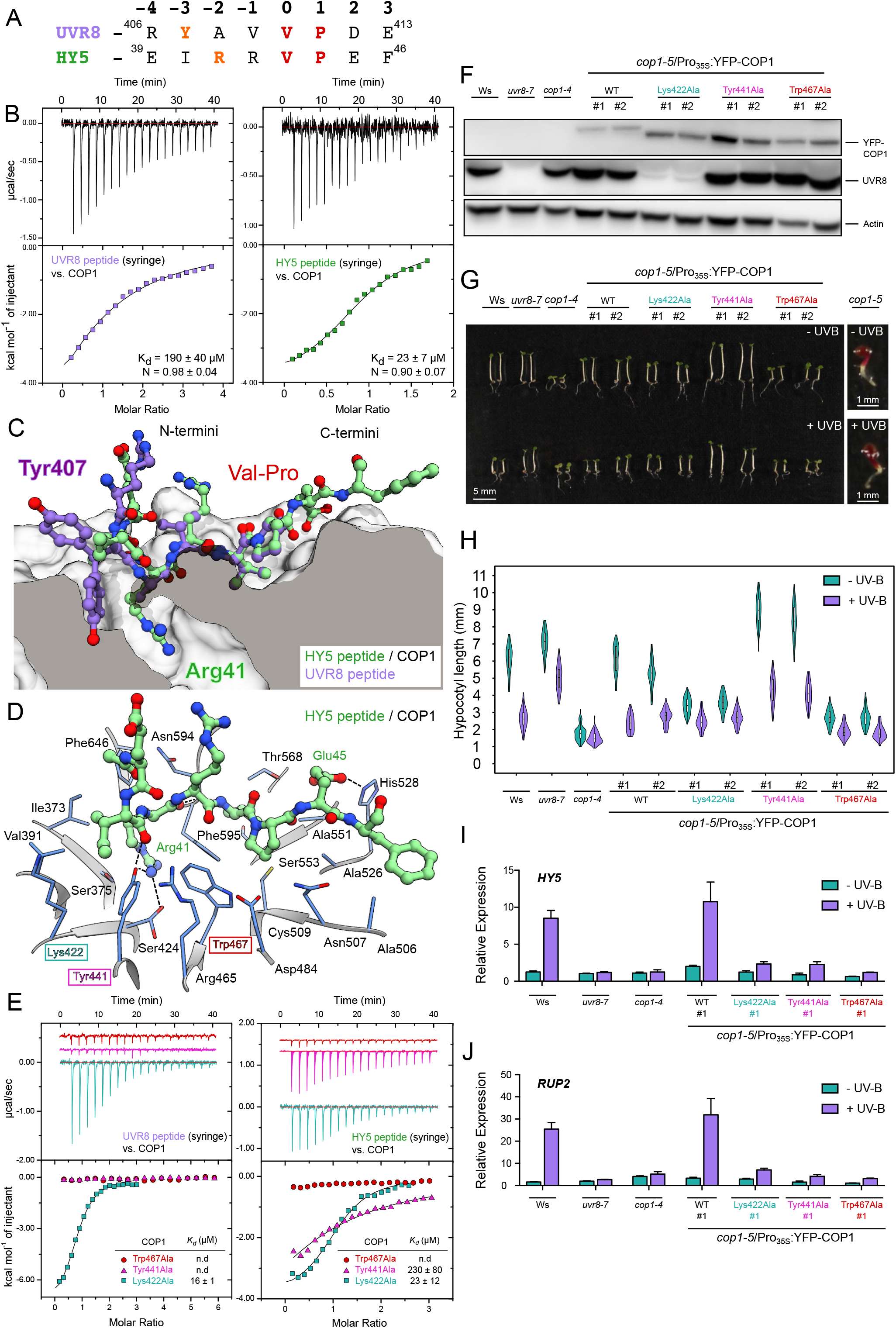
The Arabidopsis COP1 WD40 domain binds peptides representing the core VP motifs of UVR8 and HY5 with different affinities. (A) Alignment of the UVR8 and HY5 VP-peptide motifs. The conserved VP pairs are highlighted in red, and the anchor residues in orange. (B) ITC assay of the UVR8 and HY5 VP-peptides versus the COP1 WD40 domain. The top panel represents the heats detected during each injection. The bottom panel represents the integrated heats of each injection, fitted to a one-site binding model (solid line). The following concentrations were typically used (titrant into cell): UVR8 – COP1 (2500 μM in 175 μM); HY5 – COP1 (1500 μM in 175 μM). The insets show the dissociation constants (*K_d_*) and stoichiometries of binding (N) (± standard deviation). (C) Superposition of the X-ray crystal structures of the HY5 and UVR8 peptides in the VP-peptide binding site of the COP1 WD40 domain. COP1 is depicted in surface representation and belongs to the HY5 – COP1 complex. The HY5 peptide is depicted in green in ball-and-stick representation (with Arg41 labelled). The UVR8 peptide from the UVR8 – COP1 complex is superimposed on top in purple (with Tyr407 labelled), depicted in ball-and-stick representation. The surface of COP1 has been clipped to better visualize the anchor residue in the COP1 WD40 domain. (D) Ribbon diagram depicting the VP-binding site of COP1 (blue) bound to the HY5 peptide (green). Residues Lys422, Tyr441 and Trp467 are highlighted with a colored box in cyan, magenta and red, respectively. (E) ITC assays of the HY5 and UVR8 VP-peptides versus the different COP1 WD40 domain mutants (colors as in panel D). The following concentrations were typically used (titrant into cell): UVR8 – COP1^Lys422Ala^ (1000 μM in 100 μM); UVR8 – COP1^Tyr441Ala^ (2500 μM in 90 μM); UVR8 – COP1^Trp467Ala^ (2500 μM in 90 μM); HY5 – COP1^Lys422Ala^ (1500 μM in 175 μM); HY5 – COP1^Tyr441Ala^ (1600 μM in 138 μM); HY5 – COP1^Trp467Ala^ (1600 μM in 112 μM). The insets show the dissociation constants (*K_d_*) and stoichiometries of binding (N) (± standard deviation; n.d.: no detectable binding). (F) Immunoblot analysis of YFP-COP1, UVR8 and actin (loading control) protein levels in lines shown in panel G. Seedlings were grown for 4 days under white light. (G,H) Images of representative individuals (G) and quantification of hypocotyl lengths (H) of 4-day-old seedlings grown with or without supplemental UV-B. Violin and box plots are shown for *n* > 60 seedlings. (I, J) Quantitative real-time PCR analysis of (I) *HY5* and (J) *RUP2* expression. Four-day-old seedlings grown in white light were exposed to narrowband UV-B for 2 hours (+UV-B), or not (-UV-B). Error bars represent SEM of 3 biological replicates. (F-J) Lines used: wild-type (Ws), *uvr8-7, cop1-4*, cop1-5/Pro_35S_:YFP-COP1(WT), *cop1-5/Pro_35S_:YFP-COP1^Lys422Ala^, cop1-5/Pro_35S_:YFP-COP1^Tyr441Ala^, cop1-5/Pro_35S_:YFP-COP1^Trp467Ala^* and *cop1-5*. #1 and #2: independent transgenic lines.

In order to compare how the VP-peptide motifs from different plant light signaling components bind COP1, we quantified the interaction of the UVR8 and HY5 VP peptides with the recombinant Arabidopsis COP1^349-675^ WD40 domain (termed COP1 thereafter) using isothermal titration calorimetry (ITC). We find that both peptides bind COP1 with micromolar affinity, with HY5^39-48^ binding ~ 8 times stronger than UVR8^406-413^ (Figure 1B). Next, we solved crystal structures of the COP1 WD40 domain – VP-peptide complexes representing UVR8^406-413^ – COP1 and HY5^39-48^ – COP1 interactions to 1.3 Å resolution (Figure 1C). Structural superposition of the two complexes (r.m.s.d. is ~0.2 Å comparing 149 corresponding C_α_ atoms) reveals an overall conserved mode of VP-peptide binding (r.m.s.d is ~ 1.2 Å comparing 6 corresponding C_α_ atoms), with the central VP residues making hydrophobic interactions with COP1^Trp467^ and COP1^Phe595^ (buried surface area is ~500 Å^2^ in COP1) (Figures 1D and S1). COP1^Lys422^ and COP1^Tyr441^ form hydrogen bonds and salt bridges with either UVR8^Tyr407^ or HY5^Arg41^, both being anchored to the COP1 WD40 core (Figures 1C, 1D, and S1), as previously seen for the corresponding TRIB1 residues in the COP1 – TRIB1 peptide complex (Uljon et al., 2016). In our HY5^39-48^ – COP1 structure, an additional salt bridge is formed between HY5^Glu45^ and COP1^His528^ (Figure 1D). In the peptides, the residues surrounding the VP core adopt different conformations in UVR8 and HY5, which may rationalize their different binding affinities (Figures 1B and 1C). We tested this by mutating residues Lys422, Tyr441 and Trp467 in the VP-peptide binding pocket of COP1. Mutation of COP1^Trp467^ to alanine disrupts binding of COP1 to either UVR8 or HY5 derived peptides (Figures 1B and 1E). Mutation of COP1^Tyr441^ to alanine abolishes binding of COP1 to the UVR8 peptide and greatly reduces binding to the HY5 peptide (Figures 1B and 1E), in good agreement with our structures (Figure 1D). The COP1^Lys422Ala^ mutant binds HY5^39-48^ as wild-type, but increases the binding affinity of UVR8^406-413^ ~10-fold (Figures 1B and 1E). Interestingly, COP1^Lys422Ala^ interacts with full-length UVR8 also in the absence of UV-B in yeast two-hybrid assays, which is not detectable for wild-type COP1 (Figure S2A) (Rizzini et al., 2011). Moreover, COP1^Lys422Ala^ also interacts more strongly with the constitutively interacting UVR8^C44^ fragment (corresponding to the C-terminal UVR8 tail containing the VP motif) when compared to wild-type COP1 in yeast two-hybrid assays (Figure S2B). In contrast, COP1^Tyr441Ala^ and COP1^Trp467Ala^ show reduced interaction to both UVR8 and HY5 (Figure S2). A UVR8^406-413^ – COP1^Lys422Ala^ complex structure reveals the UVR8 VP-peptide in a different conformation, with UVR8^Tyr407^ binding at the surface of the VP-binding pocket (Figures S3A-S3E). In contrast, a structure of HY5^39-48^ – COP1^Lys422Ala^ closely resembles the wild-type complex (Figure S3F).

We next assessed the impact of COP1 VP-peptide binding pocket mutants in UV-B signaling assays *in planta*. The seedling-lethal *cop1-5* null mutant can be complemented by expression of YFP-COP1 driven by the CaMV 35S promoter. We introduced COP1 mutations into this construct and isolated transgenic lines in the *cop1-5* background. All lines expressed comparable levels of the YFP-fusion proteins and complemented the seedling lethality of *cop1-5* (Figures 1F, 1G and S4). We found that cop1-5/Pro_35S_:YFP-COP1^Trp467Ala^ and cop1-5/Pro_35S_:YFP-COP1^Lys422Ala^ transgenic lines have constitutively shorter hypocotyls when compared to wild-type or cop1-5/Pro_35S_:YFP-COP1 control plants (Figures 1G and 1H), in agreement with previous work (Holm et al., 2001), suggesting partially impaired COP1 activity. This is similar to the phenotype of *cop1-4* (Figures 1G and 1H), a weak *cop1* allele that is viable but fully impaired in UVR8-mediated UV-B signaling (McNellis et al., 1994; Oravecz et al., 2006; Favory et al., 2009). In contrast, *cop1-5/Pro_35S_:YFP-* COP1^Tyr441Ala^ showed an elongated hypocotyl phenotype when compared to wild-type (Figures 1G and 1H), suggesting enhanced COP1 activity. However, in contrast to YFP-COP1, none of the YFP-COP1^Lys422Ala^, YFP-COP1^Tyr441Ala^ or YFP-COP1^Trp467Ala^ restored UV-B-induced marker gene activation like *HY5, RUP2, ELIP2* and *CHS* to wild-type level (Figures 1I, 1J and S4A). Surprisingly, however, the YFP-COP1^Lys422Ala^ line showed strongly reduced UVR8 levels (Figure 1F), despite showing normal *UVR8* transcript levels (Figure S5), precluding any conclusion of the mutation’s effect on UV-B signaling per se. In contrast, YFP-COP1^Tyr441Ala^ and YFP-COP1^Trp467Ala^ were impaired in UV-B signaling, despite showing wild-type UVR8 protein levels (Figure 1F). This indicates strongly reduced UVR8 signaling, in agreement with the reduced affinity of the COP1 mutant proteins vs. UVR8^406-413^ *in vitro* (Figure 1E). Together, our crystallographic, quantitative biochemical and functional assays suggest that UVR8 and HY5 can specifically interact with the COP1 WD40 domain using sequence-divergent VP motifs, and that mutations in the COP1 VP-binding site can modulate these interactions and impair UVR8 signaling.

### High-affinity, cooperative binding of photoactivated UVR8

HY5 levels are stabilized in a UVR8-dependent manner under UV-B light (Favory et al., 2009; Huang et al., 2013). We hypothesized that COP1 is inactivated under UV-B light, by activated UVR8 preventing HY5 from interacting with COP1. Our analysis of the isolated VP-peptide motifs of UVR8 and HY5 suggests that UVR8 cannot efficiently compete with HY5 for COP1 binding. However, it has been previously found that the UVR8 β-propeller core can interact with the COP1 WD40 domain independent of its VP motif (Yin et al., 2015). We thus quantified the interaction of UV-B activated full-length UVR8 with the COP1 WD40 domain. Recombinant UVR8 expressed in insect cells was purified to homogeneity, monomerized under UV-B, and analyzed in ITC and grating-coupled interferometry (GCI) binding assays. We found that UV-B-activated full-length UVR8 binds COP1 with a dissociation constant (K_*d*_) of ~150 nM in both quantitative assays (Figures 2A and 2B) and ~10 times stronger than non-photoactivated UVR8 (Figure S6A). This 1,000 fold increase in binding affinity compared to the UVR8^406-413^ peptide indicates cooperative binding of the UVR8 β-propeller core and the VP-peptide motif. In line with this, UV-B-activated UVR8 monomers interact with the COP1 WD40 domain in analytical size-exclusion chromatography experiments, while the non-activated UVR8 dimer shows no interaction in this assay (Figure S7A).

**Figure 2:**
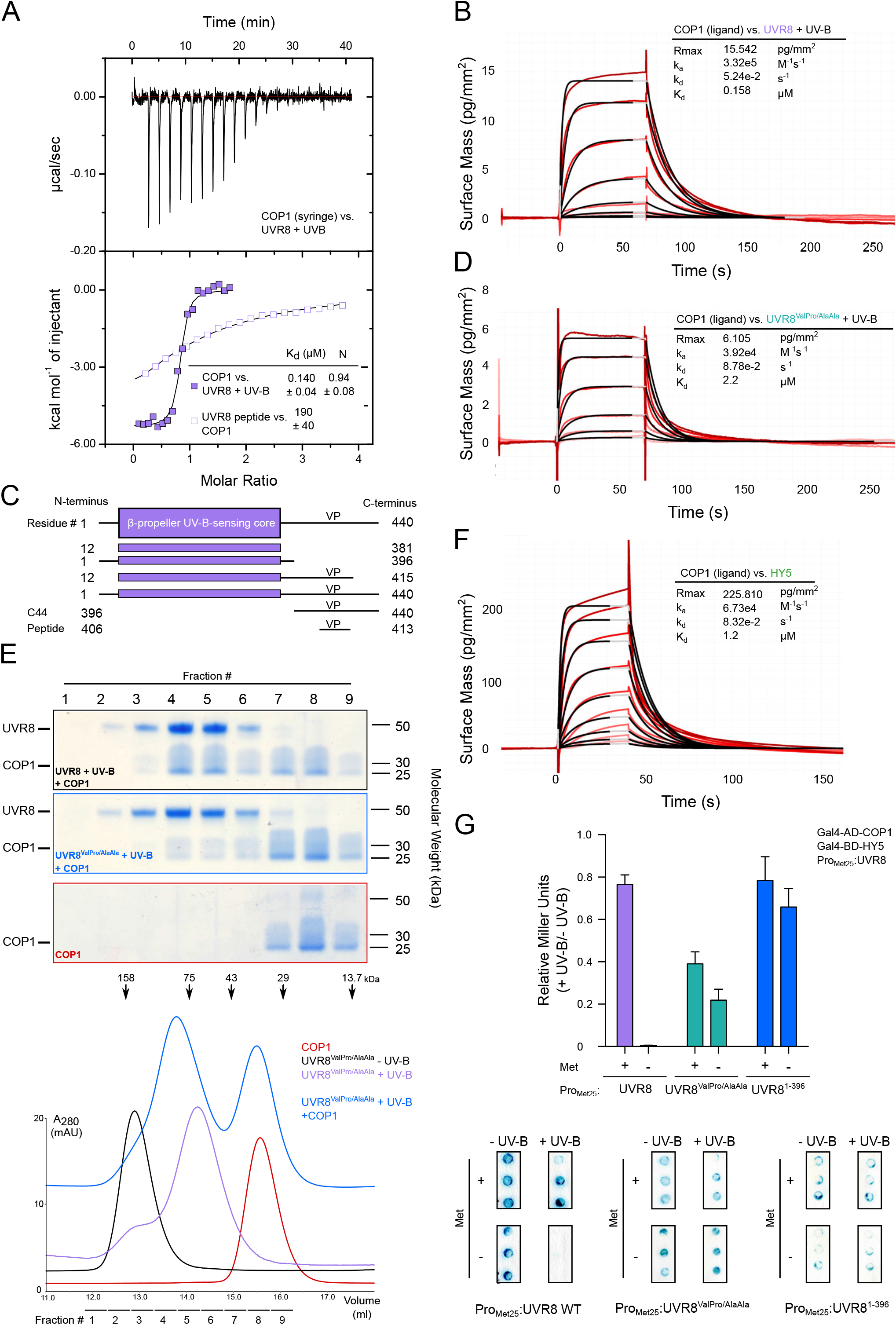
High-affinity co-operative binding of activated full-length UVR8 to the COP1 WD40 domain is mediated by its UV-B-activated β-propeller core and its C-terminal VP-peptide motif. (A) ITC assay between the COP1 WD40 domain and full-length UVR8 pre-monomerized by UV-B light. Integrated heats are shown in solid, purple squares. For comparison, an ITC experiment between UVR8 VP peptide and the COP1 WD40 domain (from Figure 1B) is also shown in open, purple squares. The following concentrations were typically used (titrant into cell): COP1 – UVR8 +UV-B (130 μM in 20 μM). The inset shows the dissociation constant (*K_d_*), stoichiometry of binding (N) (± standard deviation). (B) Binding kinetics of UVR8 pre-monomerized by UV-B versus the COP1 WD40 domain obtained by grating-coupled interferometry (GCI). Sensorgrams of UVR8 injected are shown in red, with their respective 1:1 binding model fits in black. The following amounts were typically used: ligand – COP1 (2000 pg/mm^2^); analyte – UVR8 +UV-B (highest concentration 2 μM). *k_a_* = association rate constant, *k_d_* = dissociation rate constant, *K_d_* = dissociation constant. (C) The domain organization of Arabidopsis UVR8. It consists of a UV-B-sensing β-propeller core (residues 12-381) and a long C-terminus containing the VP motif. Constructs and peptides used and their residue endings are indicated. (D) Binding kinetics of UVR8^ValPro/AlaAla^ pre-monomerized by UV-B versus the COP1 WD40 domain obtained by GCI experiments. Sensorgrams of UVR8 injected are shown in red, with their respective 1:1 binding model fits in black. The following amounts were typically used: ligand - COP1 (2000 pg/mm^2^); analyte – UVR8^ValPro/AlaAla^ +UV-B (highest concentration 2 μM). *k* = association rate constant, *k_d_* = disassociation rate constant, *K_d_* = dissociation constant. (E) Coomassie-stained SDS-PAGE gels from a size-exclusion chromatography binding assay between COP1 and UVR8^ValPro/AlaAla^ pre-monomerized by UV-B. Purified monomeric UVR8^ValPro/AlaAla^ ~ 50 kDa, COP1 WD40 a smear ~25-40 kDa. Four μM of each protein were loaded independently or mixed together. Indicated fractions were taken each of the size-exclusion chromatography runs and separated on a 10% SDS-PAGE gel. (F) Binding kinetics of HY5 versus the COP1 WD40 domain obtained by GCI experiments. Sensorgrams of HY5 injected are shown in red, with their respective 1:1 binding model fits in black. The following amounts were typically used: ligand – COP1 (2000 pg/mm^2^); analyte – HY5 (highest concentration 2 μM). *k_a_* = association rate constant, *k_d_* = dissociation rate constant, *K_d_* = dissociation constant. (G) Yeast 3-hybrid analysis of the COP1 – HY5 interaction in the presence of UVR8. (Top) Normalized Miller Units were calculated as a ratio of β-galactosidase activity in yeast grown under UV-B (+ UV-B) versus yeast grown without UV-B (-UV-B). Additionally, normalized Miller Units are reported separately here for yeast grown on media without or with 1 mM methionine, corresponding to induction (− Met) or repression (+ Met) of *Met25* promoter-driven UVR8 expression, respectively. Means and SEM for 3 biological repetitions are shown. (Bottom) representative filter lift assays. AD, activation domain; BD, DNA binding domain; Met, methionine.

As the interaction of full-length UVR8 is markedly stronger than the isolated UVR8 VP-peptide, we next dissected the individual contributions of the individual UVR8 domains to COP1 binding (Figure 2C). We find that the UV-B activated UVR8 β-propeller core (UVR8^12-381^) binds COP1 with a K_*d*_ of ~0.5 μM and interacts with the COP1 WD40 domain in size-exclusion chromatography experiments (Figures S6B and S7B). The interaction is strengthened when the C-terminus is extended to include the VP-peptide motif (UVR8^12-415^) (Figures S6B and S6C). Mutation of the UVR8 VP-peptide motif to alanines results in ~20 fold reduced binding affinity when compared to the wild-type protein (Figures 2D). However, the mutant photoreceptor is still able to form complexes with the COP1 WD40 domain in size exclusion chromatography assays (Figures 2E). We could not detect sufficient binding enthalpies to monitor the binding of UVR8^ValPro/AlaAla^ to COP1 in ITC assays nor detectable signal in GCI experiments in the absence of UV-B (Figure S8). The COP1^Lys422Ala^ mutant binds UV-B-activated full-length UVR8 with wild-type affinity, while COP1^Trp467Ala^ binds ~5 times more weakly (Figures S9A and S9B). Mutations targeting both COP1 and the UVR8 C-terminal VP-peptide motif decreases their binding affinity even further (Figure S9C). Thus full-length UVR8 uses both its β-propeller photoreceptor core and its C-terminal VP-peptide to cooperatively bind the COP1 WD40 domain when activated by UV-B light.

We next asked if UV-B-activated full-length UVR8 could compete with HY5 for binding to COP1. We produced the full-length HY5 protein in insect cells and found that it binds the COP1 WD40 domain with a K_*d*_ of ~1 μM in GCI assays (Figure 2F). For comparison, the isolated HY5 VP-peptide binds COP1 with a K_*d*_ of ~20 μM (Figure 1B). This would indicate that only the UV-B-activated UVR8 and not ground-state UVR8 (K_*d*_ ~ 150 nM vs ~ 1 μM, see above) can efficiently compete with HY5 for COP1 binding. We tested this hypothesis in yeast 3-hybrid experiments. We confirmed that HY5 interacts with COP1 in the absence of UVR8 and that this interaction is specifically abolished in the presence of UVR8 and UV-B light (Figure 2G). We conclude that UV-B-activated UVR8 efficiently competes with HY5 for COP1 binding in yeast cells, thereby impairing the COP1 – HY5 interaction under UV-B. The UVR8^ValPro/AlaAla^ and UVR8^1-396^ mutants cannot interfere with the COP1 – HY5 interaction in yeast cells (Figure 2G), suggesting that a functional UVR8 VP-peptide motif is required to compete off HY5 from COP1, in agreement with our biochemical assays.

### UVR8 – VP peptide chimeras trigger UV-B signaling *in planta*

Our findings suggest that UVR8 requires both its UV-B-sensing core and its VP-peptide motif for high affinity COP1 binding and that the UVR8 VP-peptide can inhibit the interaction of HY5 with COP1 (Figures 1 and 2) (Yin et al., 2015). This led us to speculate that any VP-peptide with sufficient binding affinity for COP1 could functionally replace the endogenous VP motif in the UVR8 C-terminus *in vivo*. We generated chimeric proteins in which the UVR8 core domain is fused to VP-containing sequences from plant and human COP1 substrates, namely HY5 and TRIB1 (Figure 3A). Arabidopsis *uvr8-7* null mutants expressing these chimeric proteins show complementation of the hypocotyl and anthocyanin phenotypes under UV-B, suggesting that all tested UVR8 chimeras are functional (Figures 3B-D, and S10). Early UV-B marker genes are also up-regulated in the lines after UV-B exposure, demonstrating that these UVR8 chimeras are functional photoreceptors, although to different levels (Figure 3E). In line with this, the UVR8^HY5C44^ chimera can displace HY5 from COP1 in yeast 3-hybrid assays (Figure 3F), can bind COP1 affinities comparable to wild-type (Figures 3G and S10) and are dimers *in vitro* that monomerize under UV-B (Figure 3H). Together, these experiments reinforce the notion that divergent VP-peptide motifs compete with each other for binding to the COP1 WD40 domain.

**Figure 3:**
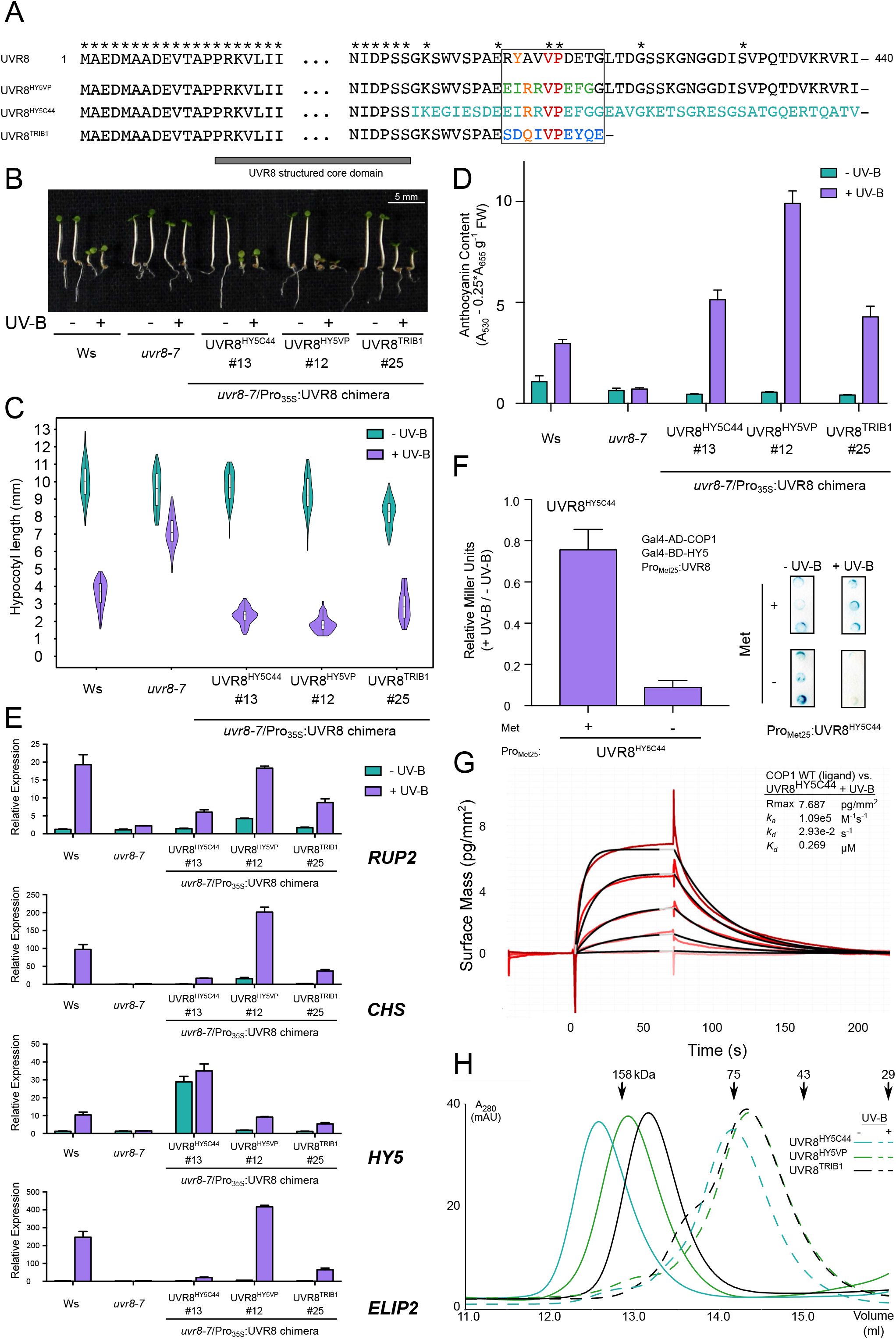
Chimeras of the UV-B-sensing UVR8 core and various VP motifs are functional in the UV-B signaling pathway. (A) Sequence alignment of the N- and C-termini of UVR8, chimera UVR8^HY5VP^ (replacing the UVR8 VP motif with the corresponding sequence from HY5), chimera UVR8^HY5C44^ (replacing the C44 domain of UVR8 with the corresponding sequence from HY5), and chimera UVR8^TRIB1VP^ (replacing the UVR8 VP motif with the TRIB1 VP motif and a truncation of the rest of the UVR8 C-terminus). The black box indicates the core VP motif of UVR8. The VP is colored in red, the anchor residues in orange, the divergent residues of the HY5 VP core sequence in green, the HY5 sequence replacing the UVR8 C-terminal 44 amino acids (C44) in cyan, and the divergent residues of the TRIB1 VP core sequence in blue. Asterisks represent amino acids identical in all constructs, amino acids 21-390 of UVR8 are not shown. The previously crystallized UVR8 core domain (PDB: 4D9S) is highlighted with a gray bar. (B) Representative image showing the phenotype of wild-type (Ws), *uvr8-7* and *uvr8-7/Pro_35S_:UVR8^HY5C44^, uvr8-7/Pro_35S_:UVR8^HY5VP^* and *uvr8-7/Pro_35S_:UVR8^TRIB1^* seedlings grown for 4 days under white light (− UV-B) or white light supplemented with UV-B (+ UV-B). (C) Quantification of hypocotyl length data shown in B. Violin and box plots are shown for *n* > 60 seedlings. (D) Anthocyanin accumulation in seedlings shown in B. Average and SEM are shown (n = 3). (E) Quantitative real-time PCR analysis of *RUP2, CHS, HY5* and *ELIP2* expression in wild-type (Ws), *uvr8-7* and *uvr8-7/Pro_35S_:UVR8^HY5C44^, uvr8-7/Pro_35S_:UVR8^HY5VP^* and *uvr8-7/Pro_35S_:UVR8^TRIB1^* seedlings in response to 2h of UV-B (+UV-B vs. −UV-B). Error bars represent SEM of 3 biological replicates. Note: the primers used to detect the *HY5* transcript abundance also bind to an identical region present in UVR8^HY5C44^ chimera. (F) Yeast 3-hybrid analysis of the COP1-HY5 interaction in the presence of the UVR8^HY5C44^ chimera. (Left pane) Normalized Miller Units were calculated as a ratio of β-galactosidase activity in yeast grown under UV-B (+ UV-B) versus yeast grown without UV-B (− UV-B). Additionally, normalized Miller Units are reported separately here for yeast grown on media without or with 1 mM methionine, corresponding to induction (Met) or repression (+ Met) of *Met25* promoter-driven UVR8 expression, respectively. Means and SEM for 3 biological repetitions are shown. (Right panel) Representative filter lift assays of the yeast analyzed in left panel. AD, activation domain; BD, DNA binding domain; Met, methionine. (G) Binding kinetics of the UVR8^HY5C44^ chimera pre-monomerized by UV-B versus the COP1 WD40 domain obtained by GCI experiments. Sensorgrams of UVR8^HY5C44^ injected are shown in red, with their respective 1:1 binding model fits in black. The following amounts were typically used: ligand – COP1 (2000 pg/mm^2^); analyte – UVR8^HY5C44^ +UV-B (highest concentration 2 μM). *k_a_* = association rate constant, *k_d_* = disassociation rate constant, *K_d_* = dissociation constant. (H) Size-exclusion chromatography assay of recombinant chimeric proteins expressed in Sf9 insect cells in the presence and absence of UV-B.

### Sequence-divergent VP-peptide motifs are recognized by the COP1 WD40 domain

Our protein engineering experiments prompted us to map core VP-peptide motifs in other plant light signaling components, including the COP1-interacting blue-light photoreceptors CRY1 and CRY2 (Yang et al., 2000; Wang et al., 2001; Yu et al., 2007; Yang et al., 2018) and the transcription factors HYH, CO/BBX1, COL3/BBX4, SALT TOLERANCE (STO/BBX24) (Holm et al., 2002; Datta et al., 2006; Jang et al., 2008; Liu et al., 2008; Yan et al., 2011), and HFR1 (Duek et al., 2004; Yang et al., 2005; Jang et al., 2005). We mapped putative VP-motifs in all these proteins and assessed their binding affinities to the COP1 WD40 domain (Figures 4A and 4B). We could detect binding for most of the peptide motifs in ITC assays, with dissociation constants in the mid-micromolar range (Figure 4A and S11). Next, we obtained crystal structures for the different peptides bound to COP1 (1.3 – 2.0 Å resolution, see Tables 1 and 2, Figures 4E, 4F and S11) to compare their peptide binding modes (Figure 4C). We found that all peptides bind in a similar configuration with the VP forming the center of the binding site (r.m.s.d.’s between the different peptides range from ~0.3 Å to 1.5 Å, comparing 5 or 6 corresponding C_α_ atoms). Chemically diverse amino-acids (Tyr/Arg/Gln) map to the -3 and -2 position and often deeply insert into the COP1 binding cleft, acting as anchor residues (Figure 4C). This suggests that the COP1 WD40 domain has high structural plasticity, being able to accommodate sequence-divergent VP-containing peptides.

**Figure 4:**
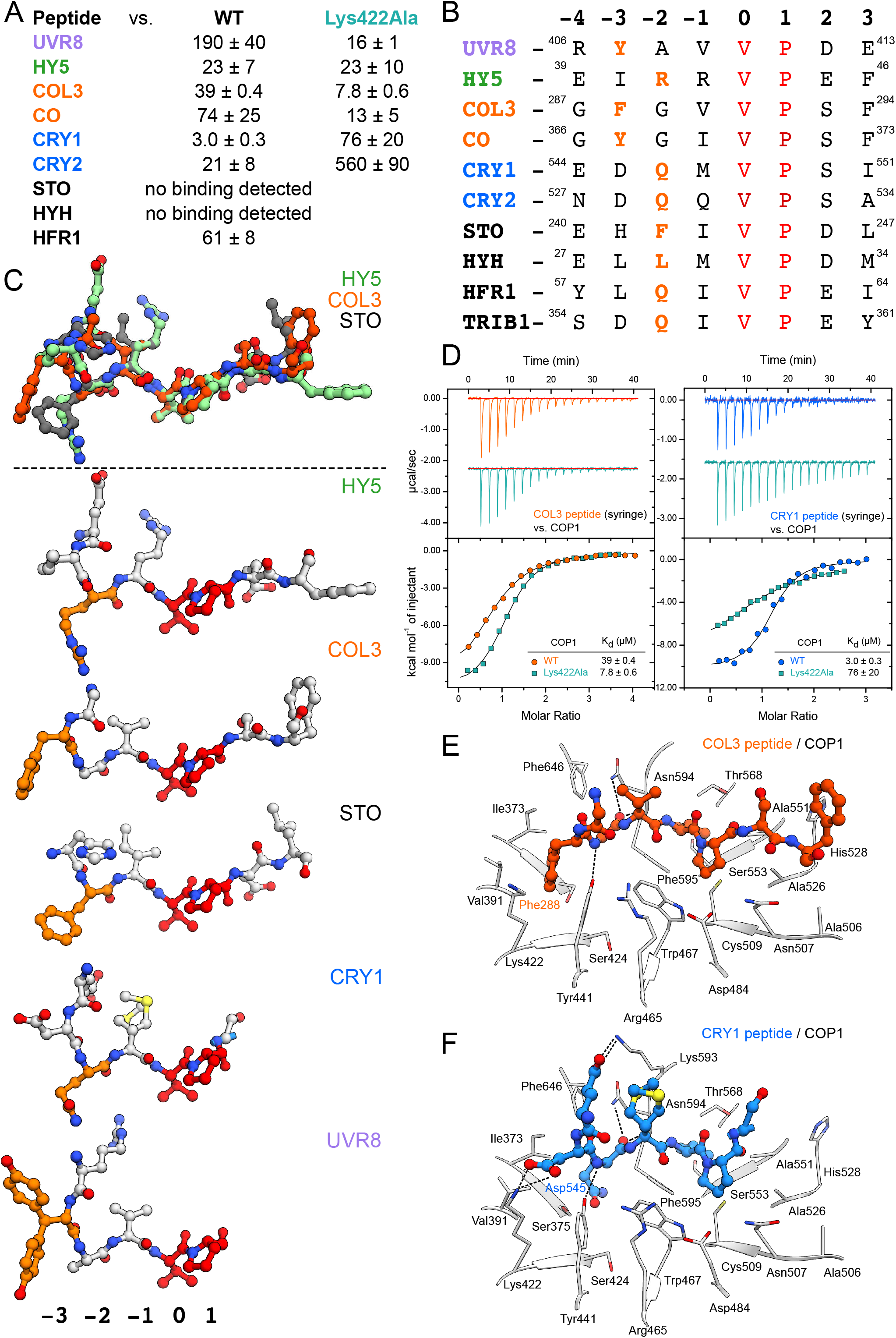
Many COP1 substrates and interacting photoreceptors contain VP-peptides that can bind the COP1 WD40 domain. (A) Table summarizing affinities (*K_d_*, dissociation constant) of VP-containing peptides versus the COP1 WD40 domain (WT) and COP1^Lys422Ala^ as determined by ITC. All values are μM (± standard deviation). (B) A sequence comparison of VP-peptide motifs that interact with the COP1 WD40 domain. The VP core is colored in red and the anchor residues in orange. The peptide sequences are numbered on a register where the V in the VP is 0. (C) (Top panel) Superposition of the VP-peptide binding modes of the HY5 (green), COL3 (orange), and STO (gray) peptides depicted in ball-and-stick representation when bound to the COP1 WD40 domain as seen in their respective x-ray crystal structures. (Bottom panel) A comparison of the HY5, COL3, STO, CRY1, and UVR8 peptides highlight the chemically diverse anchor residues and their variant positions. (D) ITC assays between the (left, orange) COL3 VP peptide and (right, cyan) CRY1 VP peptide versus the COP1 WD40 and COP1^Lys422Ala^, with a table summarizing their corresponding affinities. The following concentrations were typically used (titrant into cell): COL3 – COP1 (2240 μM in 195 μM); COL3 – COP1^Lys422Ala^ (1500 μM in 175 μM); CRY1 – COP1 (750 μM in 75 μM); CRY1 – COP1^Lys422Ala^ (1500 μM in 175 μM). The inset shows the dissociation constant (K_d_), stoichiometry of binding (N) (± standard deviation). (E,F) The crystal structure of the (E) COL3 peptide and (F) CRY1 peptide bound to the COP1 WD40 domain. The peptides are depicted in ball-and-stick representation. Selected residues from COP1 are depicted in gray in stick representation.

**Table 1:**
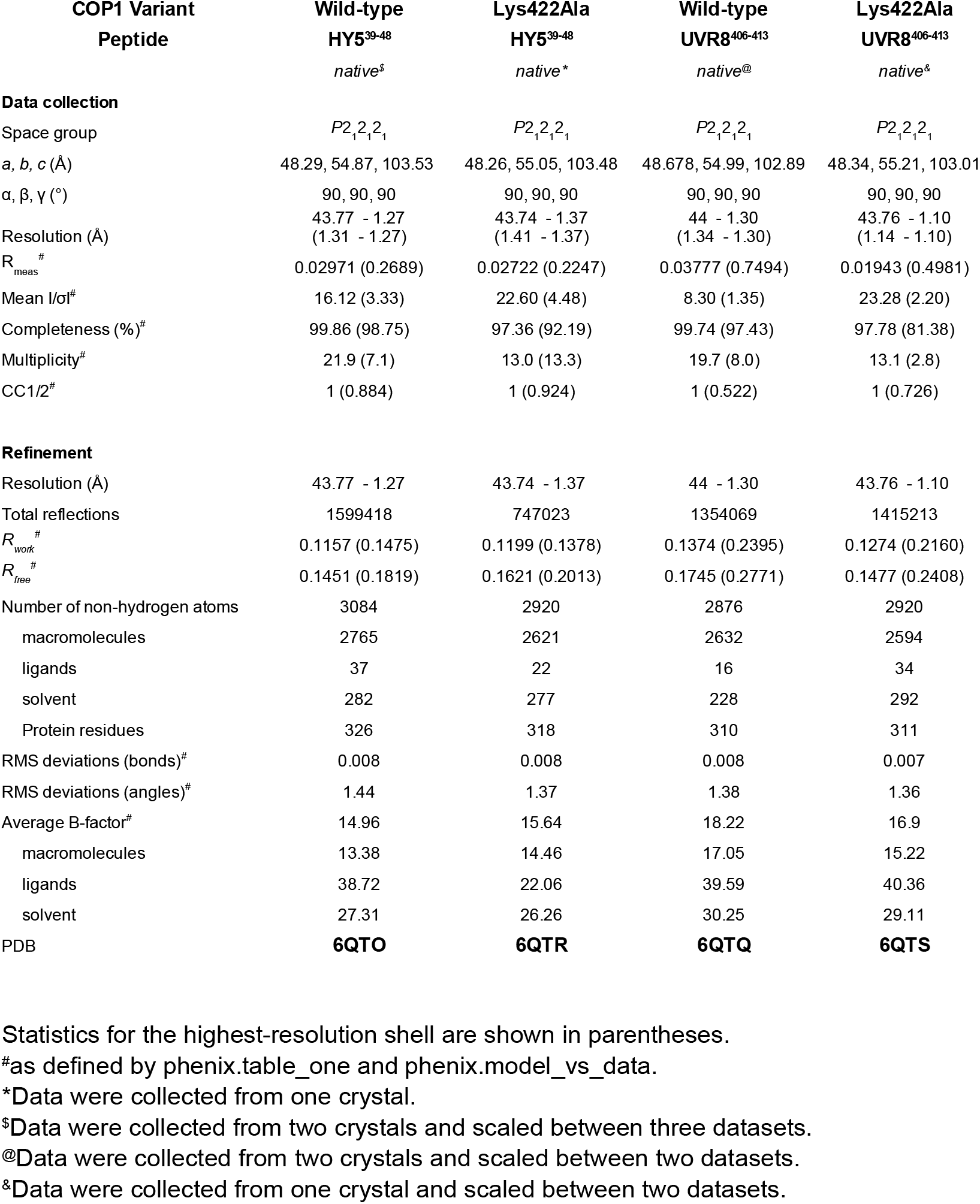
Data collection and refinement statistics

**Table 2:** Data collection and refinement statistics

| **COP1 Variant Peptide** | **Wild-type COL3<sup>287-294</sup>** | **Wild-type CRY1<sup>544-558</sup>** | **Wild-type HYH<sup>27-34</sup>** | **Wild-type HFR1<sup>57-64</sup>** | **Wild-type STO<sup>240-247</sup>** |
| --- | --- | --- | --- | --- | --- |
|  | *native\** | *native\** | *native\** | *native\** | *native\** |
| **Data collection** |  |  |  |  |  |
| Space group | $P2_12_12_1$ | $P2_12_12_1$ | $P2_12_12_1$ | $P2_12_12_1$ | $P2_12_12_1$ |
| *a*, *b*, *c* (Å) | 48.46, 55.12, 103.53 | 48.62, 55.14, 103.15 | 48.57, 55.16, 102.82 | 48.8, 55.08, 103.06 | 48.67, 54.99, 102.66 |
| $\alpha$, $\beta$, $\gamma$ (°) | 90, 90, 90 | 90, 90, 90 | 90, 90, 90 | 90, 90, 90 | 90, 90, 90 |
| Resolution (Å) | 48.66 – 1.95<br>(2.02 – 1.95) | 43.98 – 1.39<br>(1.44 – 1.39) | 43.92 – 1.51<br>(1.56 – 1.51) | 48.58 – 1.31<br>(1.36 – 1.31) | 48.48 – 1.30<br>(1.34 – 1.30) |
| $R_{\text{meas}}^{\#}$ | 0.03454 (0.2483) | 0.0561 (0.6139) | 0.05435 (0.7349) | 0.0345 (0.5801) | 0.03442 (0.746) |
| Mean $I/\sigma^{\#}$ | 24.92 (5.28) | 11.78 (1.69) | 11.65 (1.45) | 16.09 (1.82) | 16.86 (1.47) |
| Completeness (%)<sup>#</sup> | 98.21 (86.41) | 99.88 (99.04) | 99.97 (99.82) | 99.76 (97.66) | 98.68 (92.82) |
| Multiplicity<sup>#</sup> | 11.3 (4.3) | 12.7 (11.7) | 12.9 (11.6) | 12.4 (10.9) | 12.5 (11.7) |
| CC1/2<sup>#</sup> | 0.999 (0.894) | 0.999 (0.63) | 0.999 (0.595) | 1 (0.675) | 1 (0.563) |
| **Refinement** |  |  |  |  |  |
| Resolution (Å) | 48.66 – 1.95 | 43.98 – 1.39 | 43.92 – 1.51 | 48.58 – 1.31 | 48.48 – 1.30 |
| Total reflections | 233017 | 715400 | 569378 | 835400 | 845971 |
| $R_{\text{work}}^{\#}$ | 0.1389 (0.1637) | 0.1361 (0.2225) | 0.1539 (0.2620) | 0.1333 (0.2030) | 0.1413 (0.2450) |
| $R_{\text{free}}^{\#}$ | 0.1896 (0.2633) | 0.1723 (0.2765) | 0.1855 (0.2581) | 0.1760 (0.2660) | 0.1761 (0.2836) |
| Number of non-hydrogen atoms | 2745 | 2867 | 2877 | 2915 | 2891 |
| macromolecules | 2501 | 2561 | 2618 | 2619 | 2632 |
| ligands | 22 | 26 | 20 | 33 | 20 |
| solvent | 222 | 280 | 239 | 263 | 239 |
| Protein residues | 313 | 307 | 315 | 313 | 314 |
| RMS deviations (bonds)<sup>#</sup> | 0.010 | 0.008 | 0.009 | 0.008 | 0.008 |
| RMS deviations (angles)<sup>#</sup> | 1.32 | 1.36 | 1.34 | 1.34 | 1.38 |
| Average B-factor<sup>#</sup> | 16.34 | 14.74 | 20.24 | 18.19 | 17.34 |
| macromolecules | 15.56 | 13.34 | 19.28 | 16.92 | 16.27 |
| ligands | 25.25 | 24.58 | 34.24 | 25.79 | 30.65 |
| solvent | 24.30 | 26.65 | 29.55 | 29.93 | 27.99 |
| PDB | **6QTX** | **6QTW** | **6QTT** | **6QTV** | **6QTU** |

Statistics for the highest-resolution shell are shown in parentheses.
<sup>#</sup>as defined by phenix.table\_one and phenix.model\_vs\_data.
\*Data were collected from one crystal.

To experimentally investigate this property of COP1, we quantified the interaction of different VP peptides with our COP1^Lys422Ala^ mutant protein. As for UVR8 (Figures 1B and 1E), COP1^Lys422Ala^ showed increased binding affinity for some peptides such as those representing the COL3^287-294^ and CO^366-373^ VP-motifs, while it reduced binding to others, such as to CRY1^544-552^ and CRY2^527-535^ (Figures 4A and 4D). These observations may be rationalized by an enlarged VP-binding pocket in the COP1^Lys422Ala^ mutant, increasing accessibility for the COL3^Phe288^ anchor residue, and potentially abolishing interactions with CRY1^Asp545^ (Figures 4E and 4F). In yeast 3-hybrid assays we find that, similar to HY5 (Figure 2G), UV-B-activated UVR8 can efficiently compete with HYH, an N-terminal fragment of HFR1 and the CCT domain of CRY1 for binding to COP1 (Figure S12). Taken together, VP-peptide motifs of cryptochrome photoreceptors and diverse COP1 transcription factor targets all bind to the COP1 WD40 domain and UVR8 is able to compete with COP1 partners for binding.

### CRY2 and CONSTANS compete for COP1 binding

The structural plasticity of the COP1 WD40 domain is illustrated by the variable modes of binding for sequence-divergent VP motifs found in different plant light signaling components. The COP1^Lys422Ala^ mutation can modulate the interaction with different VP-peptides (Figure 4A). We noted that the cop1-5/Pro_35S_:YFP-COP1^Lys422Ala^ but not other COP1 mutants show delayed flowering when grown in long days (Figures 5A-D). This phenotype has been previously associated with mutant plants that lack the COP1 substrate CO (Figure 5B-D) (Putterill et al., 1995; Jang et al., 2008; Liu et al., 2008).

**Figure 5:**
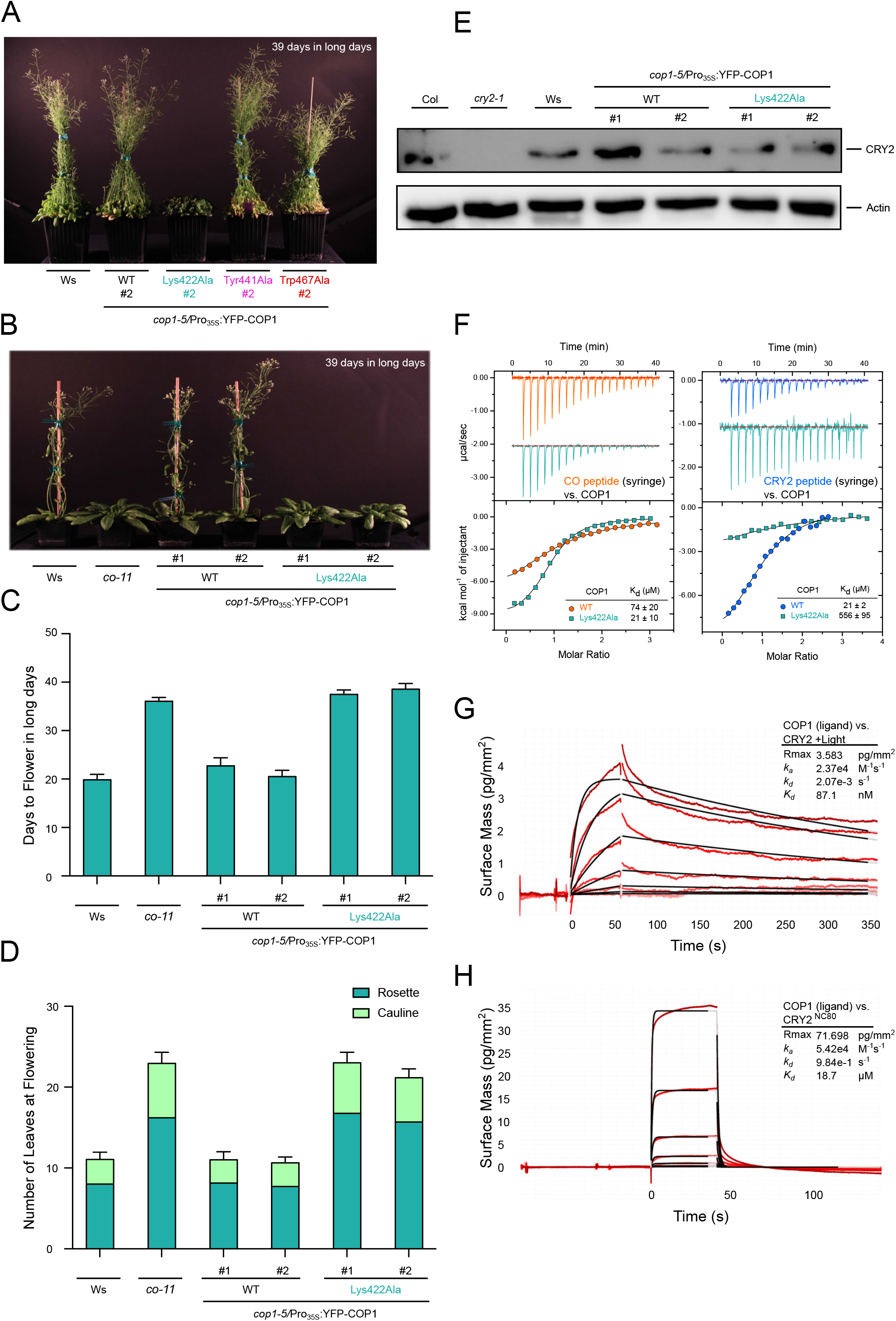
COP1^Lys422Ala^ shows a delayed flowering phenotype under long days and suggests that CO - CRY2 may compete for the COP1 VP-binding site. (A) Representative image of wild-type (Ws), cop1-5/Pro_35S_:YFP-COP1, *cop1-5/Pro_35S_:YFP-COP1^Lys422Ala^, cop1-5/Pro_35S_:YFP-COP1^Tyr441Ala^* and *cop1-5/Pro_35S_:YFP-COP1^Trp467Ala^* transgenic lines grown for 39 days in long day conditions. (B) Representative image of individual wild-type (Ws), *co-11, cop1-5/Pro_35S_:YFP-COP1* and *cop1-5/Pro_35S_:YFP-COP1^Lys422Ala^* plants grown for 39 days in long days conditions. (C) Quantification of flowering time. Means and SD are shown (*n* = 14). (D) Number of rosette and cauline leaves at flowering. Means and SD are shown (*n* = 14). (E) Immunoblot analysis of CRY2 and actin (loading control) protein levels in wild-type (Col), *cry2-1*, wild-type (Ws), *cop1-5/Pro_35S_:YFP-COP1* and *cop1-5/Pro_35S_:YFP-COP1^Lys422Ala^* seedlings grown for 4 days in darkness. (F) ITC assays between the (left, orange) CO VP-peptide and (right, cyan) CRY2 VP-peptide versus the COP1 WD40 and the COP1^Lys422Ala^ WD40 domains, with a table summarizing their corresponding affinities. The following concentrations were typically used (titrant into cell): CO – COP1 (2240 μM in 195 μM); CO – COP1^Lys422Ala^ (1200 μM in 120 μM); CRY2 – COP1 (750 μM in 70 μM); CRY2 – COP1^Lys422Ala^ (3000 μM in 175 μM). The inset shows the dissociation constant (Kd), stoichiometry of binding (N) (± standard deviation). (G,H) Binding kinetics of the (G) full-length activated cRy2 and (H) CRY2^NC80^ versus the COP1 WD40 domain obtained by GCI experiments. Sensorgrams of protein injected are shown in red, with their respective 1:1 binding model fits in black. The following amounts were typically used: ligand – COP1 (2000 pg/mm^2^); analyte – CRY2 (highest concentration 14 μM), CRY2^NC80^ (highest concentration 450 μM). *k_a_* = association rate constant, *k_d_* = dissociation rate constant.

We thus hypothesized that in COP1^Lys422Ala^ plants, binding and subsequent degradation of CO may be altered under long day conditions. *In vitro*, we found that the CO VP-peptide binds COP1^Lys422Ala^ ~4 times stronger than wild-type COP1 (Figure 5F). The same mutation in COP1 strongly reduces (~30 times) binding of the CRY2 VP-peptide *in vitro* (Figure 5F). It is of note that, in contrast to UVR8 (Figure 1F), CRY2 levels are not altered in the COP1^Lys422Ala^ background (Figure 5E).

Thus, the late flowering phenotype of the COP1^Lys422Ala^ mutant suggests that CRY2 and CO compete for COP1 binding, and that this competition is altered in the COP1^Lys422Ala^ mutant background: reduced affinity to CRY2, enhanced binding to CO – both consistent with the late flowering phenotype. In line with this, we find that recombinant light-activated full-length CRY2 binds wild-type COP1 with nanomolar affinity in quantitative GCI experiments (Figure 5G). This ~200 fold increase in binding affinity over the isolated CRY2 VP-peptide strongly suggests, that UVR8 and CRY2 both use a cooperative binding mechanism to target COP1. As a control, we tested a fragment of the CRY2 C-terminus containing the VP motif, the NC80 domain (CRY2^486-565^) (Yu et al., 2007). We found that NC80 binds COP1 with an affinity comparable to the isolated CRY2^527-535^ VP-peptide assayed by ITC (Figures 5F and 5H). Together, the COP1^Lys422Ala^ phenotypes and our biochemical assays suggest that different plant photoreceptors may use a light-induced cooperative binding mechanism, preventing COP1 from targeting downstream light signaling partners for degradation.

## DISCUSSION

The COP1 E3 ubiquitin ligase is a central hub in plant light sensing and signaling. There is strong evidence that the UV-B-sensing photoreceptor UVR8, the blue-light receptors CRY1 and CRY2 and the red/far-red discriminating phytochromes all regulate COP1 activity (Hoecker, 2017; Podolec and Ulm, 2018). The regulation of COP1 by photoreceptors enables a broad range of photomorphogenic responses, including de-etiolation, cotyledon expansion and transition to flowering, as well as UV-B light acclimation (Lau and Deng, 2012; Jenkins, 2017; Yin and Ulm, 2017; Gommers and Monte, 2018). Here we have dissected at the structural, biochemical and genetic level how the activated UVR8 and cryptochrome photoreceptors impinge on COP1 activity, by interacting with its central WD40 domain, resulting in the stabilization of COP1 substrate transcription factors. For both types of photoreceptors, interaction through a linear VP-peptide motif and a folded, light-regulated interaction domain leads to cooperative, high-affinity binding of the activated photoreceptor to COP1. We propose that in response to UV-B light, UVR8 dimers monomerize, exposing a new interaction surface that binds to the COP1 WD40 domain and releases the UVR8 C-terminal VP motif from structural restraints that prevent its interaction with COP1 in the absence of UV-B (Yin et al., 2015; Heilmann et al., 2016; Wu et al., 2019; Camacho et al., 2019). Similarly, the VP motif in the CCT domain of cryptochromes may become exposed and available for interaction upon blue-light activation of the photoreceptor (Müller and Bouly, 2015; Wang et al., 2018). Because UVR8 and CRY2 are very different in structure and domain composition, they likely use distinct interaction surfaces to target the COP1 WD40 domain, in addition to the VP-peptide motifs. The cooperative, high-affinity mode of binding enables UVR8 and cryptochromes to efficiently displace downstream signaling components such as HY5, HYH, HFR1 and CO in a light-dependent manner. Structure-guided mutations in the COP1 WD40 binding cleft resulted in the identification of the COP1^Lys422Ala^ mutant, which displays flowering phenotypes, and COP1^Tyr441Ala^ and COP1^Trp467Ala^, which display UV-B signaling phenotypes, that are all consistent with our competition model. Similar mutations have previously been shown to affect hypocotyl elongation in white light (Holm et al., 2001). Unexpectedly, COP1^Lys422Ala^ rendered the UVR8 protein unstable, preventing conclusive analysis of the effect of this COP1 mutant on UV-B signaling *in vivo*. Moreover, the mechanism behind UVR8 protein instability remains to be determined. Independent of this, it is interesting to note that the *hy4-9* mutant, which replaces the proline in the CRY1 VP-peptide motif with leucine, does not show inhibition of hypocotyl elongation under blue light (Ahmad et al., 1995). Similarly, mutations of the UVR8 VP-peptide motif or C-terminal truncations (including the *uvr8-2* allele, which has a premature stop codon at Trp400) all strongly impair UV-B signaling (Brown et al., 2005; Cloix et al., 2012; Yin et al., 2015). We now report quantitative biochemical and crystallographic analyses that reveal that UVR8 and cryptochrome photoreceptors and their downstream transcription factors all make use of VP-containing peptide motifs to target a central binding cleft in the COP1 WD40 domain. VP-containing peptides were previously identified based upon a core signature motif E-S-D-E-x-x-x-V-P-[E/D]-Φ-G, where Φ designated a hydrophobic residue (Holm et al., 2001; Uljon et al., 2016). Our structural analyses of a diverse set of VP-containing peptides now reveal that COP1 has evolved a highly plastic VP-binding pocket, which enables sequence-divergent VP motifs from different plant light signaling components to compete with each other for COP1 binding. It is reasonable to assume that many more *bona fide* VP-motifs may exist and our structures now provide sequence fingerprints to enable their bioinformatic discovery.

Interestingly, although we predict that at least some of our COP1 mutant variants (e.g. Trp467Ala) completely disrupt the interaction with VP-motif harboring COP1 targets, all COP1 variants can complement the *cop1-5* seedling lethal phenotype and largely the *cop* phenotype in darkness (Holm et al., 2001; and this work). This could imply that a significant part of COP1 activity is independent from the VP-mediated destabilization of photomorphogenesis-promoting transcription factors. It has been recently suggested that part of the *cop1* phenotype could be explained by COP1-mediated stabilization of PIFs (Pham et al., 2018). Our COP1 lines could be used to gain further insight into this aspect of COP1 activity.

Human COP1 prefers to bind phosphorylated substrates and their post-translational regulation may also be relevant in plants (Hardtke et al., 2000; Uljon et al., 2016). In this respect it is noteworthy that the full-length COP1 protein may exist as an oligomer as well as in complex with other light signaling proteins, such as SPA proteins (Seo et al., 2003; Huang et al., 2013; Sheerin et al., 2015; Holtkotte et al., 2017). The four SPA protein family members share a similar domain architecture with COP1, consisting of an N-terminal kinase-like domain, a central coiled-coil domain and a C-terminal WD40 domain (~ 45 % amino-acid identity with the COP1 WD40 domain) and are partially redundant in their activities (Yang and Wang, 2006; Ordoñez-Herrera et al., 2015). Mutations in the SPA1 WD40 domain residues Lys767 and Trp812, which correspond to COP1 residues Lys422 and Trp467, cannot complement the *spa1-3* mutant (Yang and Wang, 2006). These higher-order complexes are known to be part of some but not all light signaling pathways and could thus encode additional determinants for signaling specificity (Hoecker, 2017; Podolec and Ulm, 2018). In addition to the competition mechanism presented here, it has been observed that active cryptochrome and phytochrome receptors directly interact with SPA proteins and thereby separate COP1 from SPA proteins, which results in COP1 inactivation (Lian et al., 2011; Liu et al., 2011; Zuo et al., 2011; Lu et al., 2015; Sheerin et al., 2015). However, early UVR8 signaling is independent of SPA proteins (Oravecz et al., 2006), and may thus rely exclusively on the competition mechanism described here. For cryptochrome signaling, the VP-mediated competition and COP1-SPA disruption mechanisms are obviously not mutually exclusive but likely function in parallel *in vivo* to reinforce COP1-SPA E3 ligase inactivation in blue light signaling. Reconstitution of a photoreceptor - COP1/SPA signaling complex may offer new insights into these different targeting mechanism in the future.

## STAR METHODS CONTACT FOR REAGENT AND RESOURCE SHARING

Further information and requests for resources and reagents should be directed to, and will be fulfilled by the Lead Contact, Michael Hothorn

## EXPERIMENTAL MODEL AND SUBJECT DETAILS

### Sf9 Cell Culture

*Spodoptera frugiperda* Sf9 cells (Thermofisher) were cultured in Sf-4 Baculo Express insect cell medium (Bioconcept, Switzerland).

### Yeast Strains

The following *Saccharomyces cerevisiae* reporter strains were used: L40 (*MAT**a** trp1 leu2 his3 ade2 LYS2::lexA-HIS3 URA3::lexA-lacZ GAL4*) (Vojtek and Hollenberg, 1995), Y190 (*MAT**a** ura3-52 his3-200 lys2-801 ade2-101 trp1-901 leu2-3 112 gal4Δ gal80Δ cyh^r^2 LYS2::GAL1_UAS_-HIS3_TATA_-HIS3 MEL1 URA3::GAL1UAS-GAL1τATA-lacZ*), Y187 (*MAT**a** ura3-52 his3-200 ade2-101 trp1-901 leu2-3 112 gal4Δ met^−^ gal80Δ URA3::GAL1_UAS_-GAL1_TATA_-lacZ MEL1*) (Yeast Protocols Handbook, Clontech).

### Plants

*cop1-4* (Oravecz et al., 2006) *cop1-5* (McNellis et al., 1994), *cop1-5/Pro_35S_:YFP-COP1, cop1-5/Pro35S:YFP-COP1^Lys422Ala^, cop1-5/Pro35S:YFP-COP1^Tyr441Ala^, cop1-5/Pro35S:YFP-COP1^Trp467Ala^* (this work), *uvr8-7* (Favory et al., 2009), *uvr8-7/Pro_35S_:UVR8^HY5C44^*, *uvr8-7/Pro_35S_:UVR8^HY5VP^*, and *uvr8-7/Pro_35S_:UVR8^TRIB1^* (this work) are in the *Arabidopsis thaliana* Wassilewskija (Ws) accession. The *cry2-1* (Guo et al., 1998) mutant is in the Columbia accession. The *co-11* allele was generated in the Ws accession (this work) using CRISPR/Cas9 technology (Wang et al., 2015).

## METHOD DETAILS

### Protein expression and purification

All COP1, UVR8, HY5 and CRY2 proteins were produced as follows: The desired coding sequence was PCR amplified (see Table S1 for primers) or NcoI/NotI digested from codon-optimized genes (Geneart or Twist Biosciences) for expression in Sf9 cells. Chimeric UVR8 constructs were PCR amplified directly from vectors used for yeast 3-hybrid assays (see below). All constructs except CRY2^NC80^ were cloned into a modified pFastBac (Geneva Biotech) insect cell expression vector, via NcoI/NotI restriction enzyme sites or by Gibson assembly (Gibson et al., 2009). The modified pFastBac vector contains a tandem N-terminal His_10_-Twin-Strep-tags followed by a TEV (tobacco etch virus protease) cleavage site. CRY2^NC80^ was cloned into a modified pET-28 a (+) vector (Novagen) containing a tandem N-terminal His_10_-Twin-Strep-tags followed by a TEV (tobacco etch virus protease) cleavage site by Gibson assembly. Mutagenesis was performed using an enhanced plasmid mutagenesis protocol (Liu and Naismith, 2008).

pFastBac constructs were transformed into DH10MultiBac cells (Geneva Biotech), white colonies indicating successful recombination were selected and bacmids were purified by the alkaline lysis method. Sf9 cells were transfected with the desired bacmid with Profectin (AB Vector). eYFP-positive cells were observed after 1 week and subjected to one round of viral amplification. Amplified, untitred P2 virus (between 5 – 10 % culture volume) was used to infect Sf9 cells at a density between 1-2 x 10^6^ cells/mL. Cells were incubated for 72 h at 28°C before the cell pellet was harvested by centrifugation at 2000 x g for 20 minutes and stored at −20°C.

CRY2^NC80^ was produced in transformed Rosetta (DE3) pLysS cells. *E. coli* were grown in 2xYT broth. 1 L of broth was inoculated with 20 mL of a saturated overnight preculture, grown at 37°C until OD600 ~ 0.5, induced with IPTG at a final concentration of 0.2 mM, and then shaken for another 16 h at 18°C. The cell pellet was harvested by centrifugation at 2000 x g for 20 minutes and stored at −20°C.

Every 1 L of Sf9 or bacterial cell culture was dissolved in 25 mL of Buffer A (300 mM NaCl, 20 mM HEPES 7.4, 2 mM BME), supplemented with glycerol (10% v/v), 5 μL Turbonuclease and 1 Roche cOmplete Protease inhibitor tablets. Dissolved pellets were lysed by sonication and insoluble materials were separated by centrifugation at 60000 x g for 1 h at 4°C. The supernatant was filtered through tandem 1 μm and 0.45 μm filters before Ni^2+^-affinity purification (HisTrap excel, GE Healthcare). Ni^2+^-bound proteins were washed with Buffer A and eluted directly into a coupled Strep-Tactin Superflow XT column (IBA) by Buffer B (500 mM NaCl, 500 mM imidazole pH 7.4, 20 mM HEPES pH 7.4). Twin-Strep-tagged-bound proteins on the Strep-Tactin column were washed with Buffer A and eluted with 1X Buffer BXT (IBA). Proteins were cleaved overnight at 4°C with TEV protease. Cleaved proteins were subsequently purified from the protease and affinity tag by a second Ni^2+^-affinity column or by gel filtration on a Superdex 200 Increase 10/300 GL column (GE Healthcare). Proteins were concentrated to 3 – 10 mg/mL and either used immediately or aliquoted and frozen directly at −80°C. Typical purifications were from 2 to 5 L of cell pellet.

All protein concentrations were measured by absorption at 280 nm and calculated from their molar extinction coefficients. Molecular weights of all proteins were confirmed by MALDI-TOF mass spectrometry. SDS-PAGE gels to assess protein purity are shown in Figure S13.

For UVR8 monomerization and activation by UV-B, proteins were diluted to their final assay concentrations (as indicated in the figure legends) in Eppendorf tubes and exposed to 60 minutes at max intensity (69 mA) under UV-B LEDs (Roithner Lasertechnik GmbH) on ice.

### Analytical size-exclusion chromatography

Gel filtration experiments were performed using a Superdex 200 Increase 10/300 GL column (GE Healthcare) pre-equilibrated in 150 mM NaCl, 20 mM HEPES 7.4, 2 mM BME. 500 μl of the respective protein solution or a mixture (~4 μM per protein) was loaded sequentially onto the column and elution at 0.75 ml/min was monitored by UV absorbance at 280 nm.

### Isothermal titration calorimetry (ITC)

All experiments were performed in a buffer containing 150 mM NaCl, 20 mM HEPES 7.4, 2 mM BME. Peptides were synthesized and delivered as lyophilized powder (Peptide Specialty Labs GmbH) and dissolved directly in buffer. The peptides were centrifuged at 14000 x g for 10 minutes and only the supernatant was used. The dissolved peptide concentrations were calculated based upon their absorbance at 280 nm and their corresponding molar extinction coefficient. Typical experiments consisted of titrations of 20 injections of 2 μL of titrant (peptides) into the cell containing COP1 at a 10-fold lower concentration. Typical concentrations for the titrant were between 500 and 3000 μM for experiments depending on the affinity. Experiments were performed at 25°C and a stirring speed of 1000 rpm on an ITC200 instrument (GE Healthcare). All data were processed using Origin 7.0 and fit to a one-site binding model after background buffer subtraction.

### Grating-coupled interferometry (GCI)

The Creoptix WAVE system (Creoptix AG), a label-free surface biosensor was used to perform GCI experiments. All experiments were performed on 2PCH or 4PCH WAVEchips (quasi-planar polycarboxylate surface; Creoptix AG). After a borate buffer conditioning (100 mM sodium borate pH 9.0, 1 M NaCl; Xantec) COP1 (ligand) was immobilized on the chip surface using standard amine-coupling: 7min activation (1:1 mix of 400mM *N*-(3-dimethylaminopropyl)-*N*’-ethylcarbodiimide hydrochloride and 100 mM N-hydroxysuccinimide (both Xantec)), injection of COP1 (10 μg/mL) in 10 mM sodium acetate pH 5.0 (Sigma) until the desired density was reached, and final quenching with 1 M ethanolamine pH 8.0 for 7 min (Xantec). Since the analyte CRY2 showed nonspecific binding on the surface, BSA (0.5% in 10 mM sodium acetate, pH 5.0; BSA from Roche) was used to passivate the surface between the injection of COP1 and ethanolamine quenching. For a typical experiment, the analyte (UVR8/CRY2) was injected in a 1:3 dilution series (highest concentrations as indicated in the figure legends) in 150 mM NaCl, 20 mM HEPES 7.4, 2 mM BME at 25°C. Blank injections were used for double referencing and a dimethylsulfoxide (DMSO) calibration curve for bulk correction. Analysis and correction of the obtained data was performed using the Creoptix WAVEcontrol software (applied corrections: X and Y offset; DMSO calibration; double referencing) and a one-to-one binding model with bulk correction was used to fit all experiments.

### Protein crystallization and data collection

Crystals of co-complexes of HY5^39-48^ – COP1^349-675^, UVR8^406-413^ – COP1^349-675^, HY5^39-48^ – COP1^349-675, Lys422Ala^, UVR8^406-413^ – COP1^349-675, Lys422Ala^ and COL3^287-294^ – COP1^349-675^ were grown in sitting drops and appeared after several days at 20°C when 5 mg/mL of COP1 supplemented with 3 to 10 fold molar excess in peptide was mixed with two-fold (v/v) more mother liquor (1:2 ratio; protein:buffer) containing 2 M (NH_4_)_2_SO_4_ and 0.1 M HEPES pH 7.4 or 0.1 M Tris pH 8.5. Crystals were harvested and cryoprotected in mother liquor supplemented with 25% glycerol and frozen under liquid nitrogen.

Crystals of complexes of HYH^27-34^ – COP1^349-675^, HFR1^57-64^ – COP1^349-675^, STO^240-247^ – COP1^349-675^ and CRY1^544-552^ – COP1^349-675^ were grown in sitting drops and appeared after several days at 20°C when 5 mg/mL of COP1 supplemented with 3 to 10 fold molar excess in peptide was mixed with two-fold (v/v) more mother liquor (1:2 ratio; protein:buffer) containing 1.25 M sodium malonate pH 7.5. Crystals were harvested and cryoprotected in mother liquor supplemented with 25% glycerol and frozen under liquid nitrogen.

All datasets were collected at beam line PX-III of the Swiss Light Source, Villigen, Switzerland. Native datasets were collected with λ=1.03 Å. All datasets were processed with XDS (Kabsch, 1993) and scaled with AIMLESS as implemented in the CCP4 suite (Winn et al., 2011).

### Crystallographic structure solution and refinement

The structures of all the peptide – COP1 WD40 complexes were solved by molecular replacement as implemented in the program Phaser (McCoy et al., 2007), using PDB-ID 5IGO as the initial search model. The final structures were determined after iterative rounds of model-building in COOT (Emsley and Cowtan, 2004), followed by refinement in REFMAC5 (Murshudov et al., 2011) as implemented in CCP4 and phenix.refine (Adams et al., 2010). Polder omit maps were generated for the UVR8^406-413^ - COP1 structure by omitting residue Tyr407 of the bound peptide as implemented in phenix.polder. Final statistics were generated as implemented in phenix.table_one. All figures were rendered in UCSF Chimera (Pettersen et al., 2004).

### Plant transformation

To generate the cop1-5/Pro_35S_:YFP-COP1 line, COP1 cloned into pENTR207C was introduced into the Gateway-compatible binary vector pB7WGY2 (Karimi et al., 2002). COP1 mutated versions were generated by PCR-based site-directed mutagenesis, cloned into pDONR207 and then introduced in pB7WGY2 (Karimi et al., 2002). The wild-type version of the construct contains an additional Gateway-cloning related 14 amino acids linker sequence between the YFP and COP1. *cop1-5* heterozygous plants (kan^R^) were transformed using the floral dip method (Clough and Bent, 1998). Lines homozygous for the *cop1-5* mutation and for single locus insertions of the *Pro_35S_:YFP-* COP1 transgene were selected.

To generate lines expressing chimeric UVR8 receptors, the *HY5* and *TRIB1* sequences were introduced by PCR to the *UVR8* coding sequences as indicated, and the chimeras were cloned into the Gateway-compatible binary vector pB2GW7 (Karimi et al., 2002) for transformation into the *uvr8-7* mutant background. Lines homozygous with single genetic locus transgene insertions were selected.

To generate a *co* mutant in the Ws background, designated *co-11*, plants were transformed with the CRISPR/Cas9 binary vector pHEE401E (Wang et al., 2015) in which an sgRNA specific to the *CO* CDS was inserted (see Table S1). A plant was isolated in T2 and propagated, harboring a 1 base-pair insertion after the codon for residue Asp137 leading to a frameshift and a premature stop codon after four altered amino acids (*D*PRGR*; *D* representing Asp137 in CO, * representing the premature stop).

### Plant growth conditions

For experiments at seedling stage, Arabidopsis seeds were surface-sterilized and sown on halfstrength MS medium (Duchefa), stratified in the dark at 4°C for 48 h, and grown under aseptic conditions in controlled light conditions at 21°C. For hypocotyl length and anthocyanin measurements, the MS medium was supplemented with 1% sucrose (AppliChem). For flowering experiments, Arabidopsis plants were grown on soil in long day (16 h / 8 h; light / dark cycles) growth chambers at 21°C.

UV-B treatments were performed as described before, using Osram L18W/30 tubes, supplemented with narrowband UV-B from Philips TL20W/01RS tubes (Oravecz et al., 2006; Favory et al., 2009).

### Hypocotyl length assays

For hypocotyl length measurements, at least 60 seedlings were randomly chosen, aligned and scanned. Measurements were performed using the NeuronJ plugin of ImageJ (Meijering et al., 2004). Violin and box plots were generated using the ggplot2 library in R (Wickham, 2009).

### Anthocyanin quantification

Accumulation of anthocyanin pigments was assayed as described previously (Yin et al., 2012). In brief, 40 to 60 mg of seedlings were harvested, frozen and grinded before adding 250 μl acidic methanol (1% HCl). Samples were incubated on a rotary shaker for 1 hour and the supernatant was collected and absorbances at 530 and 655 nm were recorded using a spectrophotometer. Anthocyanin concentration was calculated as (A_530_ - 2.5 * A_655_) / mg, where mg is the fresh weight of the sample.

### Protein extraction and immunoblotting

For total protein extraction, plant material was grinded and incubated with an extraction buffer composed of 50 mM Na-phosphate pH 7.4, 150 mM NaCl, 10% (v/v) glycerol, 5 mM EDTA, 0.1% (v/v) Triton X-100, 1 mM DTT, 2 mM Na_3_VO_4_, 2 mM NaF, 1% (v/v) Protease Inhibitor Cocktail (Sigma) and 50 μM MG132, as previously described (Arongaus et al., 2018).

Proteins were separated by electrophoresis in 8% (w/v) SDS–polyacrylamide gels and transferred to PVDF membranes (Roth) according to the manufacturer’s instructions (iBlot dry blotting system, ThermoFisher Scientific), except for CRY2 immunoblots, which were transferred on nitrocellulose membranes (Bio-Rad).

For protein gel blot analyses, anti-UVR8^(426-440)^ (Favory et al., 2009), anti-UVR8^(1-15)^ (Yin et al., 2015), anti-UVR8^(410-424)^ (Heijde and Ulm, 2013), anti-CHS (sc-12620; Santa Cruz Biotechnology), anti-GFP (Living Colors^®^ A.v. Monoclonal Antibody, JL-8; Clontech), anti-actin (A0480; Sigma-Aldrich) and anti-CRY2^(588-602)^ (Eurogentec, raised against the peptide N’-CEGKNLEGIQDSSDQI-C’ and affinity purified) were used as primary antibodies. Horseradish peroxidase-conjugated antirabbit and anti-mouse immunoglobulins (Dako) were used as secondary antibodies. Signal detection was performed using the ECL Select Western Blotting Detection Reagent (GE Healthcare) and an Amersham Imager 680 camera system (GE Healthcare).

### Quantitative real-time PCR

RNA was extracted from seedlings using the RNeasy Plant Mini kit (Qiagen) following the manufacturer’s instructions. RNA samples were treated for 20 min with RNA-free DNAse (Qiagen) followed by addition of DEPC-treated EDTA for inactivation at 65°C for 10 min. Reverse transcription was performed using Taqman Reverse Transcription reagents (Applied Biosystems), using a 1:1 mixture of oligo dT and random hexamer primers. Quantitative real-time PCR was performed on a QuantStudio 5 Real-Time PCR system (ThermoFisher Scientific) using PowerUp SYBR Green Master Mix reagents (Applied Biosystems). Gene-specific primers for *CHS, COP1, ELIP2, HY5, RUP2*, and *UVR8* were described before (Favory et al., 2009; Gruber et al., 2010; Heijde et al., 2013) and *18S* expression was used as reference gene (Vandenbussche et al., 2014), expression values were calculated using the ΔΔCt method (Livak and Schmittgen, 2001) and normalized to the wild-type. Each reaction was performed in technical triplicates; data shown are from three biological repetitions.

### Flowering time assays

For quantitative flowering time measurements, the number of days to flowering was determined at bolting, and rosette and cauline leaf numbers were counted when the inflorescence reached approximately 1 cm in length (Möller-Steinbach et al., 2010).

### Yeast 2-hybrid and 3-hybrid assays

For yeast 2-hybrid assay, *COP1* and its mutated variants were introduced into pGADT7-GW (Marrocco et al., 2006; Yin et al., 2015) and *HY5, UVR8* and *UVR8^C44^* were introduced into pBTM116-D9-GW (Stelzl et al., 2005; Yin et al., 2015; Binkert et al., 2016). Vectors were cotransformed into the L40 strain (Vojtek and Hollenberg, 1995) using the lithium acetate-based transformation protocol (Gietz, 2014). Transformants were selected and grown on SD/-Trp/-Leu medium (Formedium). For analysis of β-galactosidase activity, enzymatic assays using chlorophenol red-β-D-galactopyranoside (Roche Applied Science) as substrate were performed as described (Yeast Protocols Handbook, Clontech).

For yeast 3-hybrid analysis, pGADT7-GW-COP1 was transformed into the Y190 strain (Harper et al., 1993). *HY5, HYH* and *HFR1^N186^* were cloned into the BamHI/EcoRI site of pBridge (Clontech), and *UVR8, UVR8^ValPro/AlaAla^, UVR8^1-396^* and *UVR8^HY5C44^* were cloned into the BglII/PstI cloning site, followed by transformation into the Y187 strain (Harper et al., 1993). Transformants were mated, selected and grown on SD/-Trp/-Leu/-Met medium (Formedium). For analysis of β-galactosidase activity, filter-lift assays were performed as described (Yeast Protocols Handbook, Clontech). Enzymatic assays using chlorophenol red-β-D-galactopyranoside (Roche Applied Science) were performed as described (Yeast Protocols Handbook, Clontech). For repression of Pro_Met25_:UVR8 expression, SD/-Trp/-Leu/-Met medium was supplemented with 1 mM L-methionine (Fisher Scientific).

For assays, yeast cells were grown for 2 days at 30°C in darkness or under narrow-band UV-B (Philips TL20W/01RS; 1.5 μmol m^−2^ s^−1^), as indicated.

## QUANTIFICATION AND STATISTICAL ANALYSIS

Data of ITC and GCI binding assays are reported with errors as indicated in their Figure legends.

## DATA AND SOFTWARE AVAILABILITY

The atomic coordinates of complexes have been deposited with the following Protein Data Bank accession codes: HY5^39-48^ – COP1^349-675^ : **6QTO**, UVR8^406-413^ – COP1^349-675^ : **6QTQ**, HY5 – COP1^349-675, Lys422Ala^: **6QTR** UVR8 – COP1^349-675, Lys422Ala^: **6QTS** HYH^27-34^ – COP1^349-675^: **6QTT** STO^240-247^ – COP1^349-675^ : **6QTU**, HFR1^57-64^ – COP1^349-675^ : **6QTV**, CRY1^544-552^ – COP1^349-675^ : **6QTW** and COL3^287-294^ – COP1^349-675^ : **6QTX**.

## Acknowledgments

This work was supported by an HHMI International Research Scholar Award to M.H., the Swiss National Science Foundation (grant number 31003A_175774 to R.U.), and the European Research Council (ERC) under the European Union’s Seventh Framework Programme (grant no. 310539 to R.U.). R.P. was supported by an iGE3 PhD Salary Award, K.L. was supported by an EMBO Longterm Fellowship (ALTF 493-2015). We thank Luis Lopez-Molina for technical assistance with the UVB LEDs.

---

**Figure S1:**
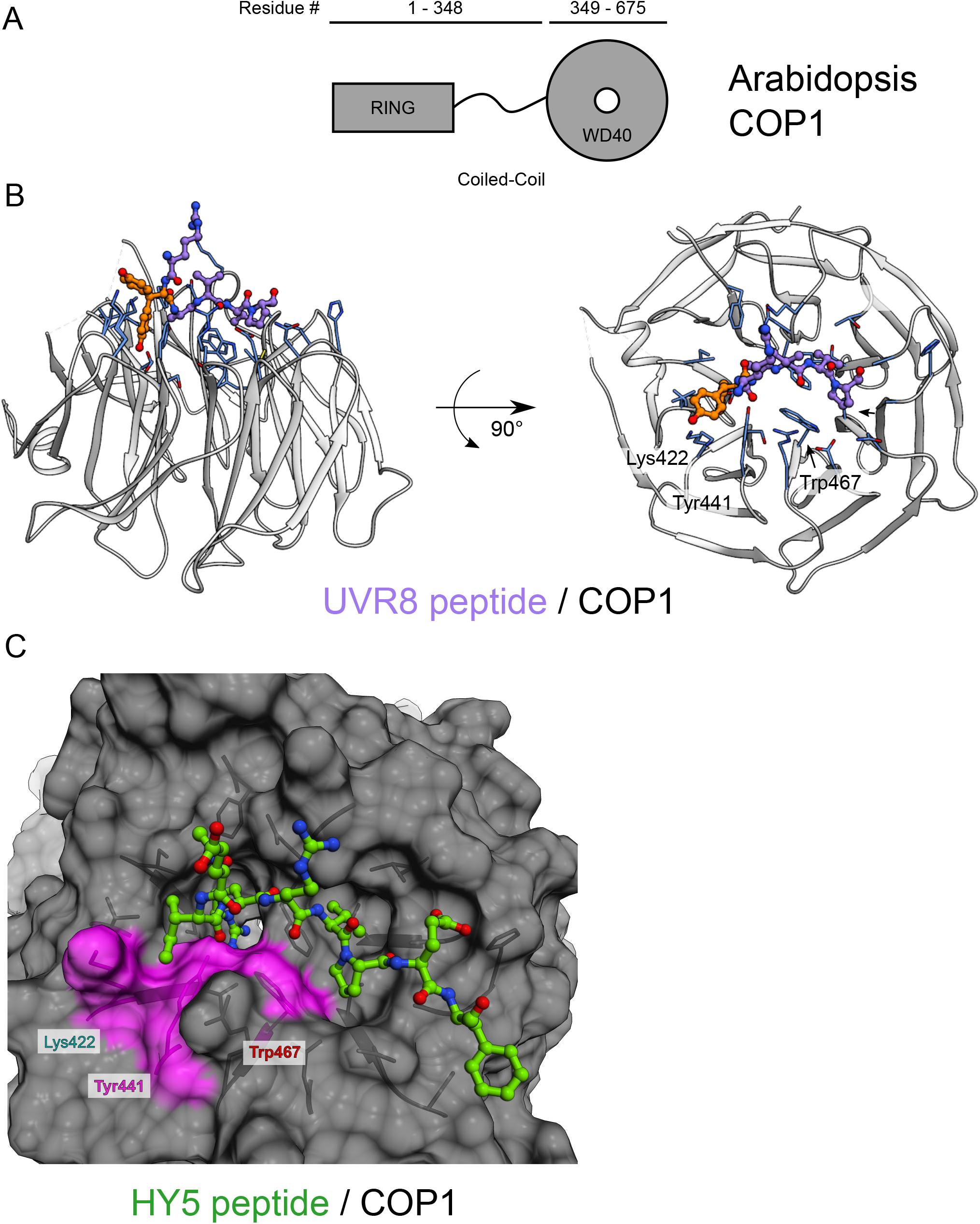
X-ray structures of the COP1 WD40 domains bound to UVR8 and HY5 VP-peptides. (A) The domain organization of the COP1 protein from Arabidopsis. It consists of an N-terminal RING domain followed by a central coiled-coil domain (residues 1-348) and a WD40 domain (residues 349-675). (B) Ribbon diagram of the COP1 WD40 domain (in gray, key residues in blue) bound to the UVR8 VP peptide (in purple). (C) A surface representation of the HY5 peptide binding site of COP1. COP1 is depicted in surface representation, the HY5 peptide is depicted in green in ball-and-stick representation. Selected residues which were mutated in this work (see Figure 1) are highlighted in magenta along with their corresponding accessible surface area.

**Figure S2:**
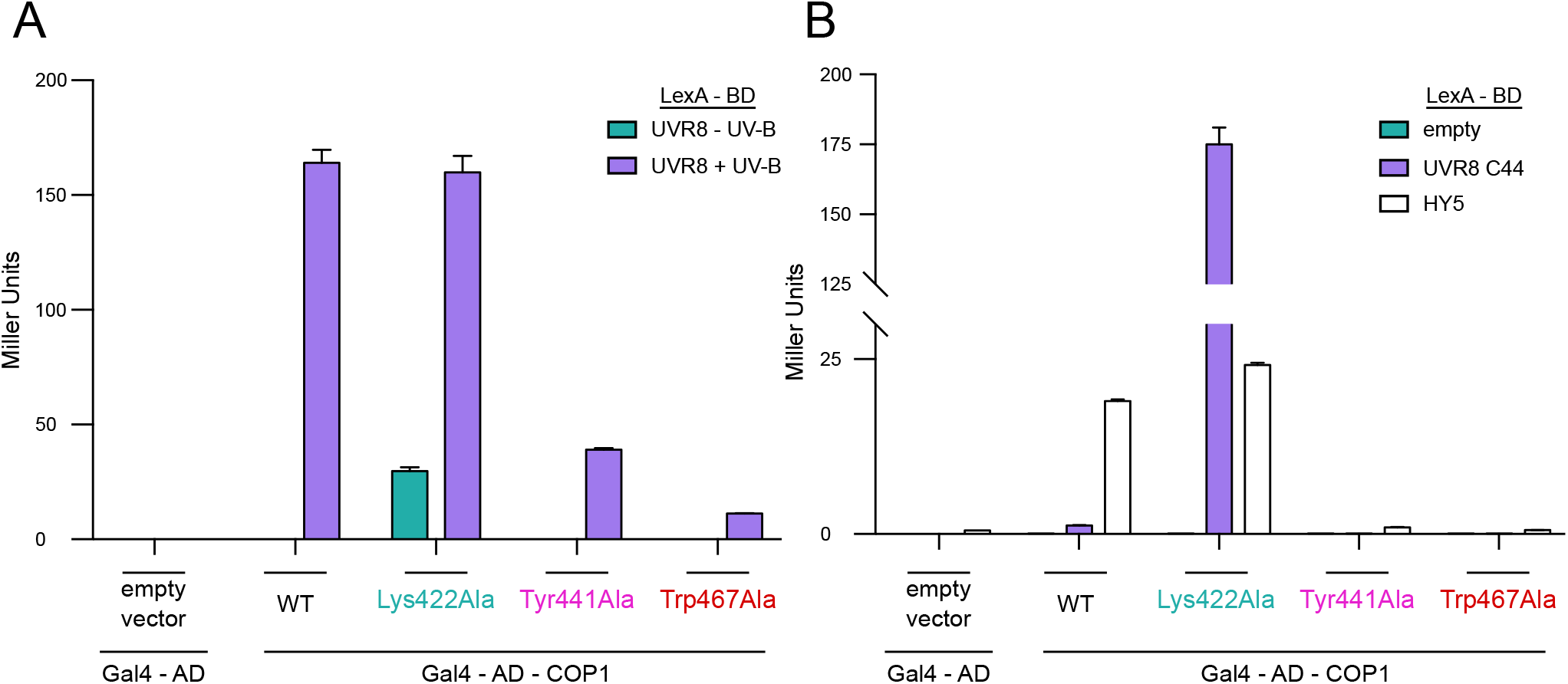
Interaction analysis of COP1 mutants with UVR8 and HY5 in yeast. (A, B) Yeast two-hybrid analysis of interactions of COP1 (WT) and COP1 mutants (COP1^Lys422Ala^, COP1^Tyr441Ala^, and COP1^Trp467Ala^) with UVR8 (+/− UV-B), UVR8^044^ and HY5. Means and SEM for 3 biological repetitions are shown. AD, activation domain; BD, DNA binding domain.

**Figure S3:**
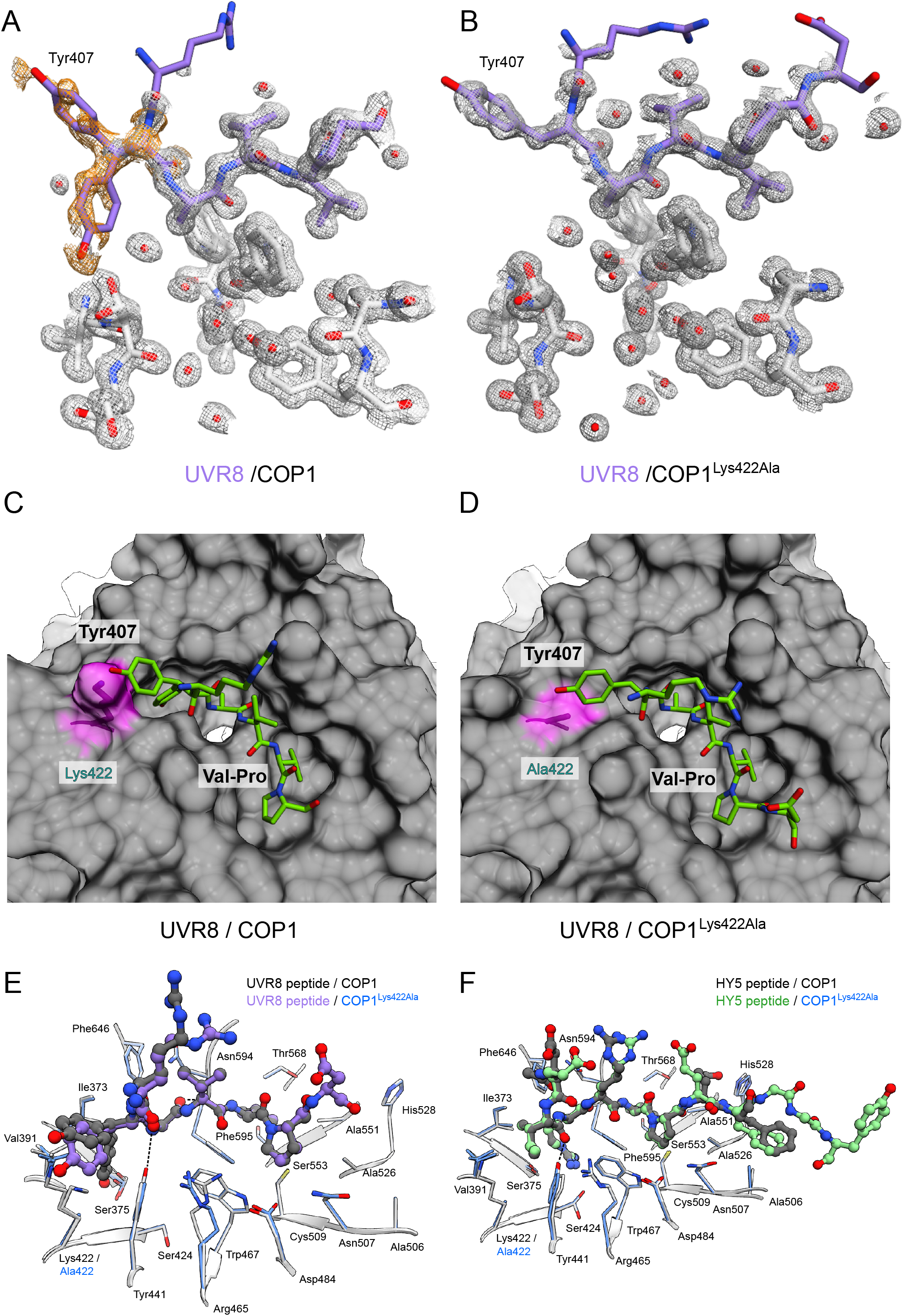
X-ray crystal structures of COP1 wild-type and COP1^Lys422Ala^ WD40 domains bound to the UVR8 VP-peptide. (A) The crystal structure of the UVR8 VP-peptide depicted in purple in stick representation bound to the COP1 WD40 domain depicted in white also in stick representation. Only selected residues and water molecules in red are shown. The white mesh represents the 2mFo-DFc electron density map contoured around all atoms depicted at a level of 1 σ. The orange mesh represents the polder omit map depicted at a level of 2.5 σ and contoured only around Tyr407 of the UVR8 VP-peptide. Two different conformers of Tyr407 were visible in the electron density and were modeled as shown. (B) The crystal structure of the UVR8 VP-peptide depicted in purple in stick representation bound to the COP1^Lys422Ala^ WD40 domain depicted in white in stick representation. Only selected residues and water molecules in red are shown. The white mesh represents the 2mFo-DFc electron density map contoured around all atoms depicted at a level of 1 σ. (C,D) A surface representation of the UVR8 VP-peptide binding site of (C) wild-type COP1 and (D) COP1^Lys422Ala^. COP1 is depicted in surface representation, the UVR8 peptide is depicted in green in ball-and-stick representation. Lys422 or Ala422 is highlighted in magenta along with their corresponding accessible surface area. (E,F) Superposition of the X-ray structures of the (E) UVR8 and (F) HY5 VP-peptides bound to the COP1 WD40 domain versus COP1^Lys422Ala^. The UVR8 VP-peptides are depicted in ball-and-stick representation. Selected residues from COP1 are depicted stick representation. The wild-type structure is gray. In the COP1^Lys422Ala^ structure, the peptide is highlighted in purple and the residues in blue.

**Figure S4:**
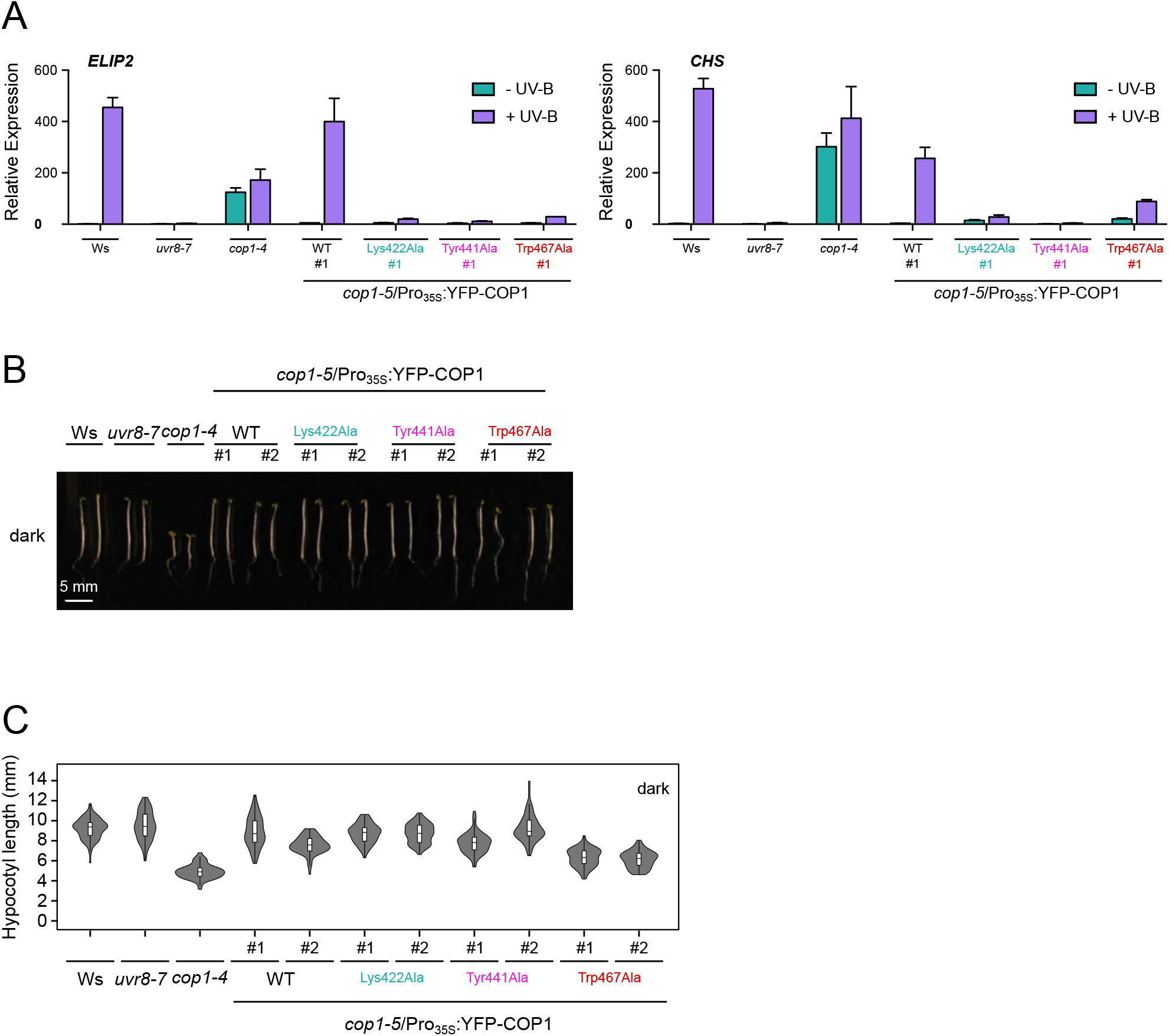
Characterization of COP1 mutant lines. (A) Quantitative real-time PCR analysis of *ELIP2* and *CHS* expression. Four-day-old seedlings grown in white light were exposed to narrowband UV-B for 2 hours (+UV-B), or not (-UV-B). Error bars represent SEM of 3 biological replicates. (B,C) Images of representative individuals (B) and quantification of hypocotyl lengths (C) of 4-day-old seedlings grown in darkness. Violin and box plots are shown for *n* > 60 seedlings. (A-C) Lines used: wild-type (Ws), *uvr8-7, cop1-4*, cop1-5/Pro_35S_:YFP-COP1 (WT), *cop1-5/Pro_35S_:YFP-COP1^Lys422Ala^*, cop1-5/Pro_35S_:YFP-*COP1^Tyr441Ala^* and *cop1-5/Pro_35S_:YFP-COP1^Trp467Ala^* seedlings. #1 and #2: independent transgenic lines.

**Figure S5:**
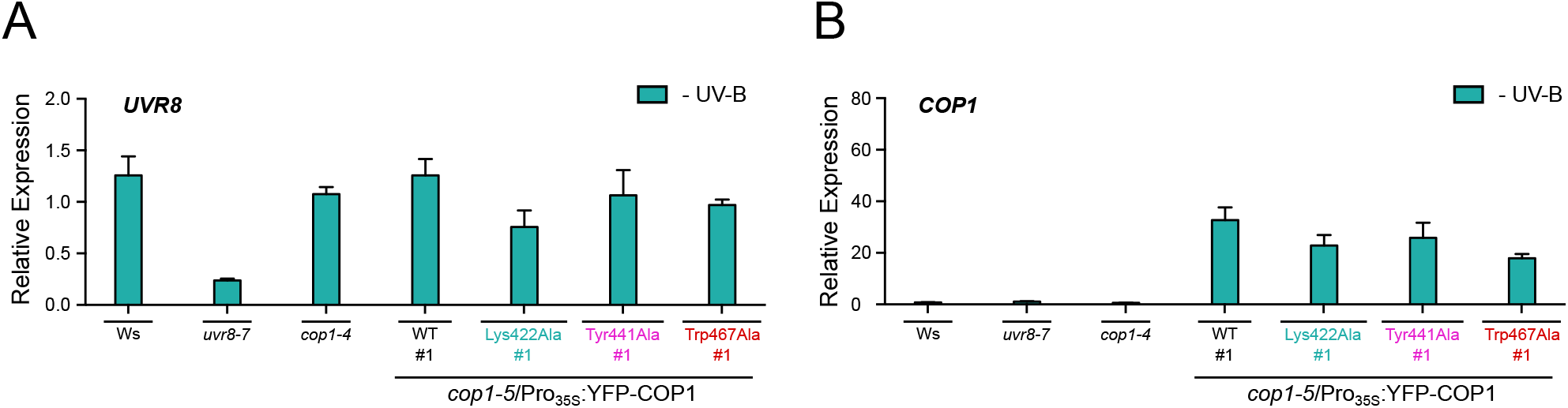
Analysis of the COP1^Lys422Ala^ mutant. (A,B) Quantitative real-time PCR analysis of (A) *UVR8* and (B) *COP1* expression in wild-type (Ws), *uvr8-7, cop1-4* and *cop1-5/Pro_35S_:YFP-COP1* (WT), *cop1-5/Pro_35S_:YFP-COP1^Lys422Ala^, cop1-5/Pro_35S_:YFP-COP1^Tyr441Ala^* and *cop1-5/Pro_35S_:YFP-COP1^Trp467Ala^* seedlings grown for 4 days under weak white light. Error bars represent SEM of 3 biological replicates.

**Figure S6:**
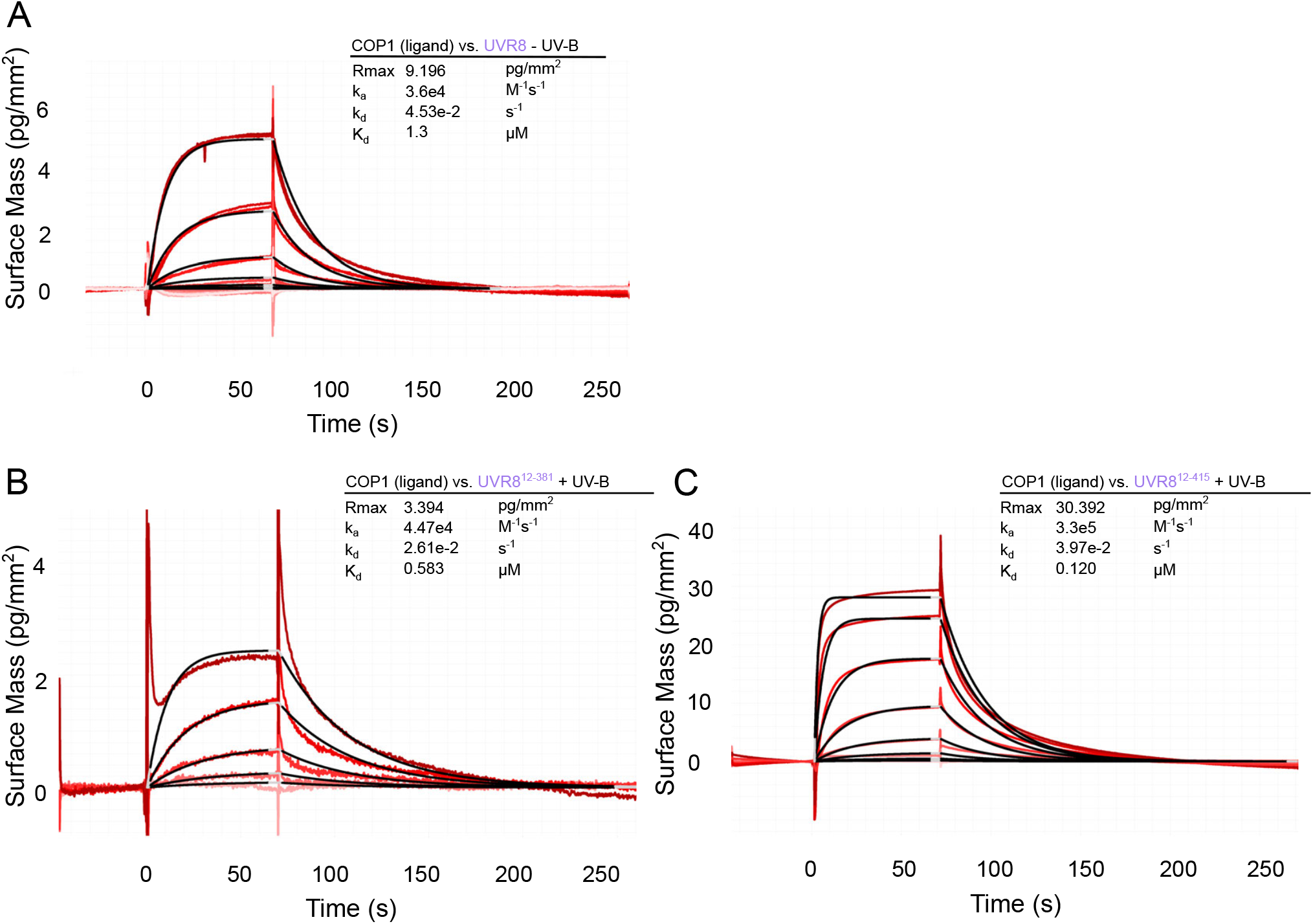
UVR8 is able to bind the COP1 WD40 domain weakly in the absence of UV-B light and the UVR8 core domain is able to bind the COP1 WD40 domain independent of the VP-peptide motif containing C-terminus in response to UV-B by GCI experiments. (A-C) Binding kinetics of (A) UVR8 in the absence of UV-B, (B) UVR8^12-381^ pre-monomerized by UV-B or (C) UVR8^12-415^ pre-monomerized by UV-B versus the COP1 WD40 domain obtained by GCI. Sensorgrams of UVR8 injected are shown in red, with their respective 1:1 binding model fits in black. The following amounts were typically used: ligand - COP1 (2000 pg/mm^2^); analyte – UVR8 and variants (highest concentration 2 μM). *k_a_* = association rate constant, *k_d_* = dissociation rate constant, *K_d_* = dissociation constant.

**Figure S7:**
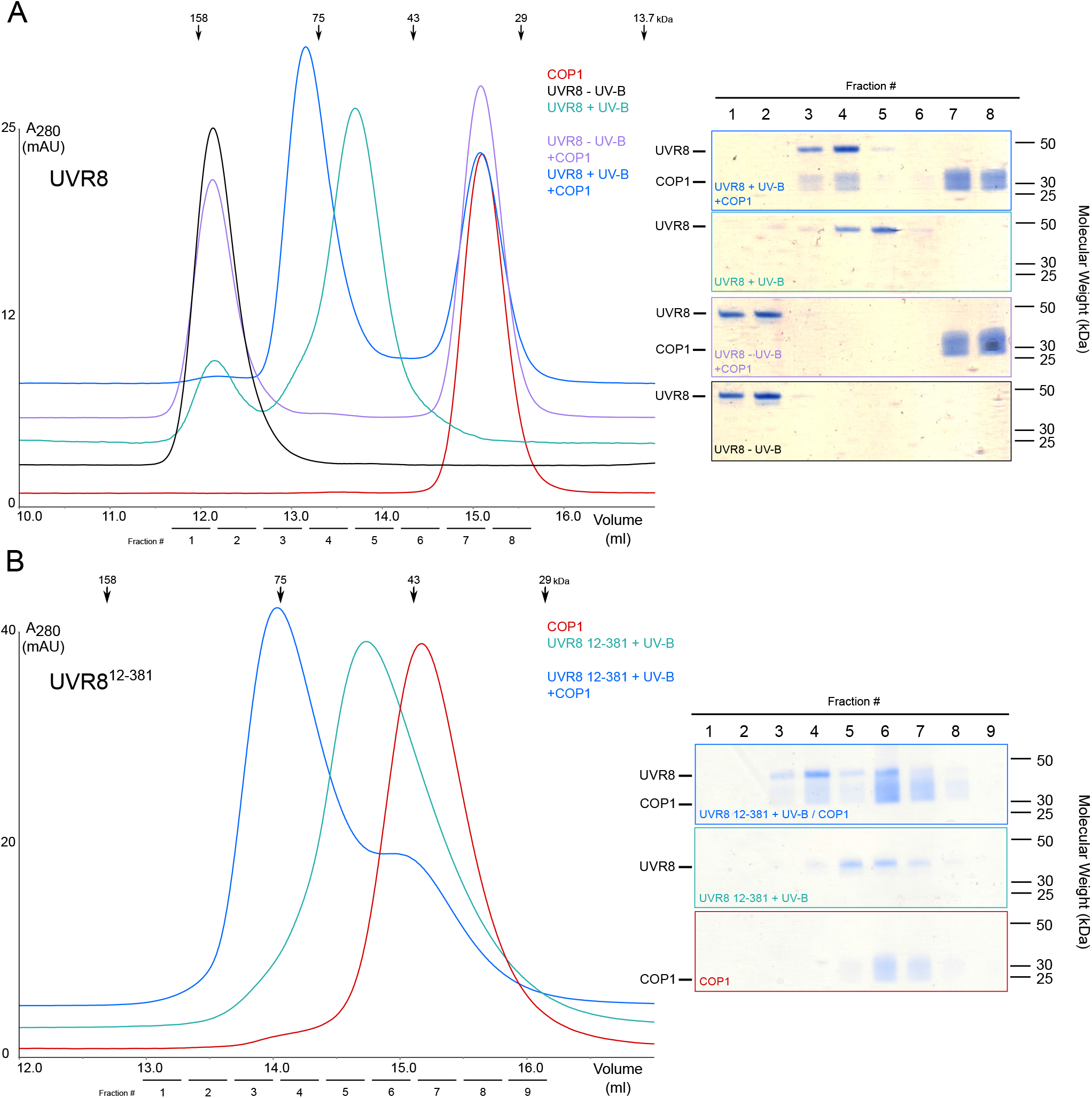
Only UV-B-activated UVR8 is able to bind COP1 in size-exclusion chromatography binding assays. (A) Coomassie-stained SDS-PAGE gels from a size-exclusion chromatography binding assay between the COP1 WD40 domain and UVR8 in the presence and absence of UV-B. Purified monomeric UVR8 ~ 50 kDa. Four μM of each protein or a mix of proteins were loaded on to a Superdex 200 Increase 10/300 GL column. Indicated fractions were taken each of the size-exclusion chromatography runs and separated on a 10 % SDS-PAGE gel. (B) Coomassie-stained SDS-PAGE gels from a size-exclusion chromatography binding assay between the COP1 WD40 domain and UVR8^12-381^ in the presence and absence of UV-B. Purified monomeric UVR8^12-381^ ~ 40 kDa. Four μM of each protein or a mix of proteins were loaded on to a Superdex 200 Increase 10/300 GL column. Indicated fractions were taken each of the size-exclusion chromatography runs and separated on a 10% SDS-PAGE gel.

**Figure S8:**
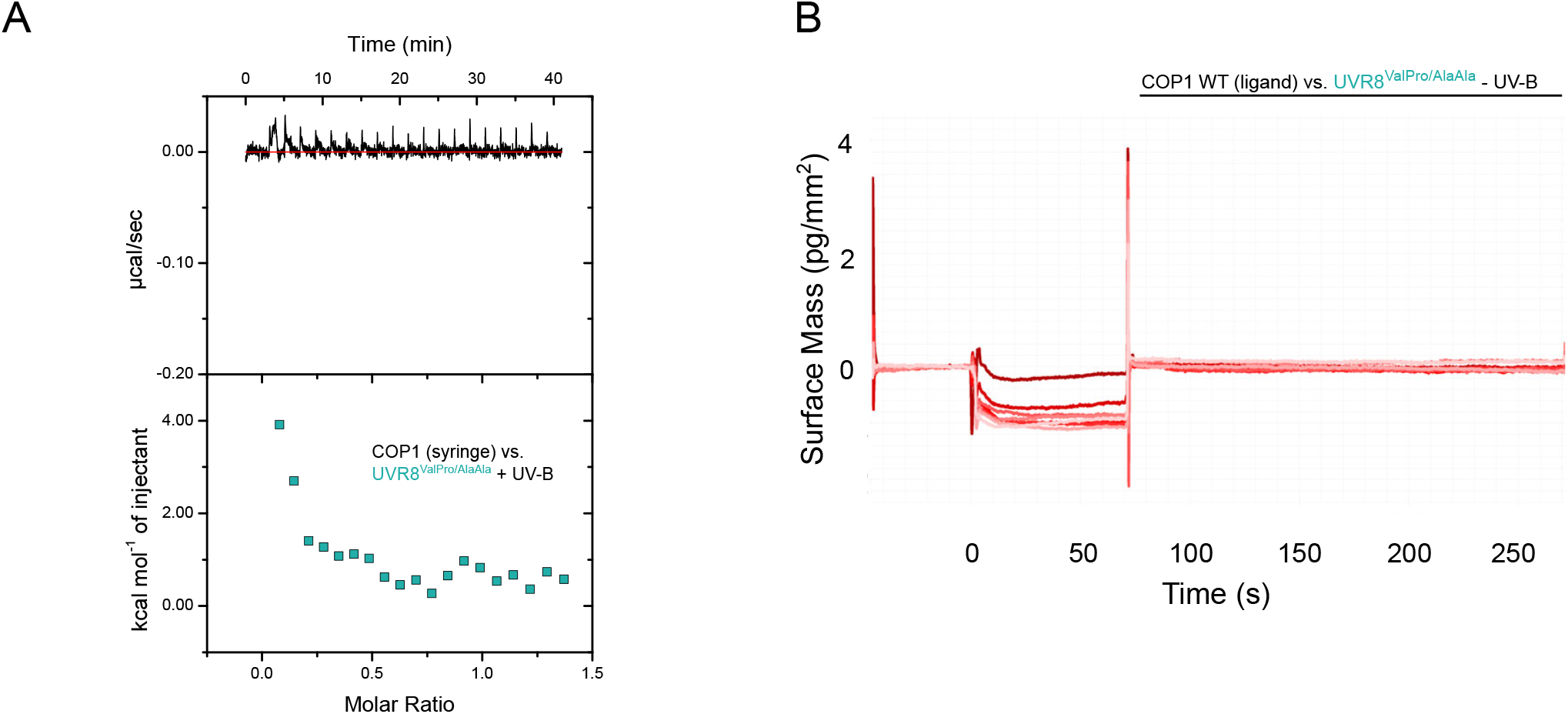
UV-B-activated UVR8^ValPro/AlaAla^ binding to the COP1 WD40 domain is not detectable by ITC experiments and UVR8^ValPro/AlaAla^ does not bind to the COP1 WD40 domain in the absence of UV-B. (A) ITC experiment between the COP1 WD40 domain and full-length UVR8^ValProAlaAla^ pre-monomerized by UV-B. Integrated heats are shown in solid, cyan squares. The following concentrations were typically used (titrant into cell): UVR8^VaPro/AlaAla^ – COP1 (130 μM in 20 μM). (B) No binding was observed for UVR8^ValPro/AlaAla^ in the absence of UV-B versus the COP1 WD40 domain obtained by GCI experiments. Sensorgrams of UVR8^ValPro/AlaAla^ injected are shown in red. The following amounts were typically used: ligand - COP1 (2000 pg/mm^2^); analyte - UVR8^ValPro/AlaAla^ +UV-B (highest concentration 2 μM). *k_a_* = association rate constant, *k_d_ =* dissociation rate constant, *K_d_* = dissociation constant.

**Figure S9:**
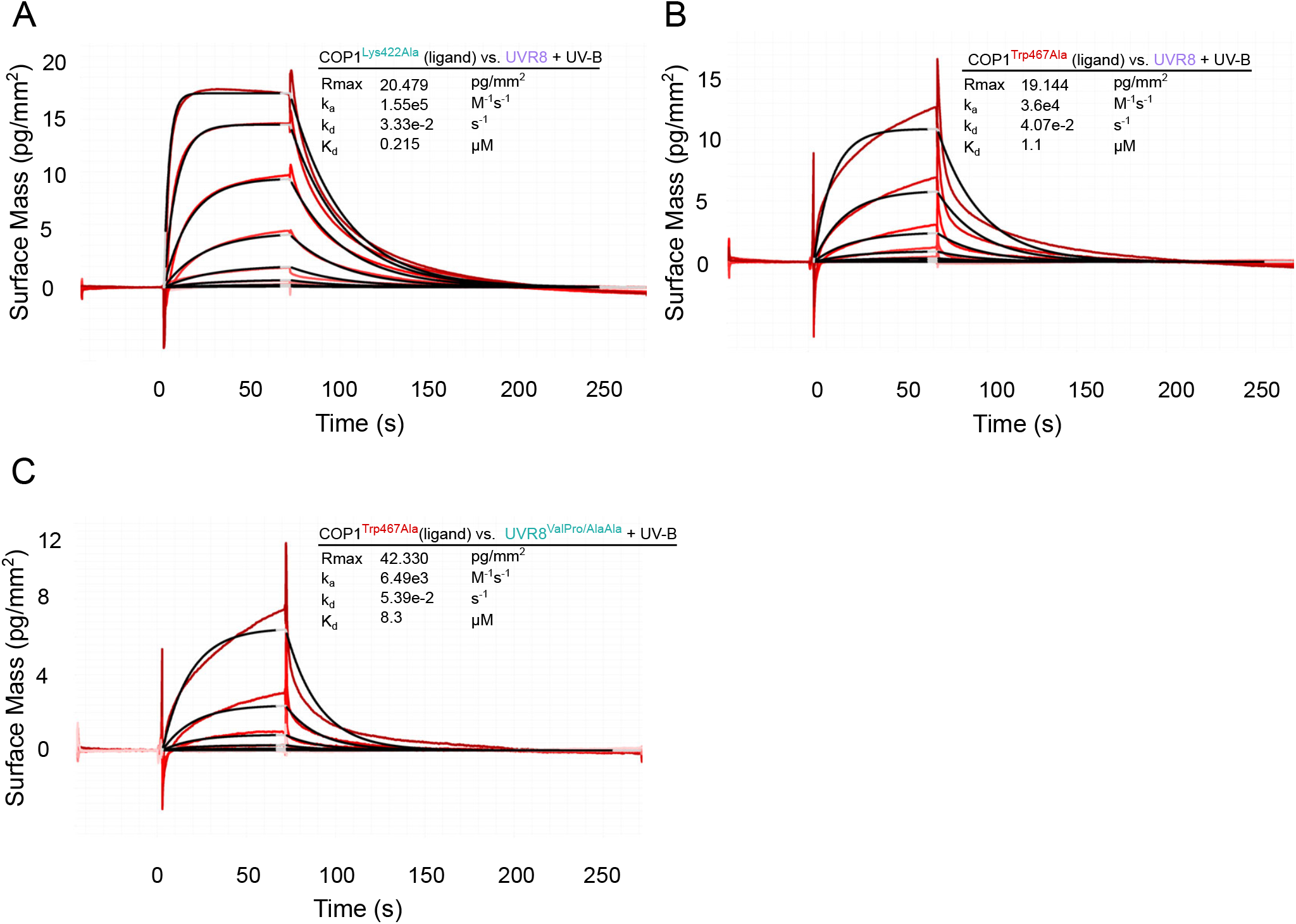
Mutations targeting the COP1 VP-binding site and the UVR8 VP motif both affect binding. (A,B) Binding kinetics of UVR8 pre-monomerized by UV-B versus the (A) COP1^Lys422Ala^ WD40 domain or (B) COP1^Trp467Ala^ WD40 domain obtained by GCI experiments. Sensorgrams of UVR8 injected are shown in red, with their respective 1:1 binding model fits in black. The following amounts were typically used: ligand – COP1^Lys422Ala^ (2000 pg/mm^2^) or – COP1^Trp467Ala^ (4000 pg/mm^2^); analyte – UVR8 +UV-B (highest concentration 2 μM). *k_a_* = association rate constant, *k_d_* = dissociation rate constant. (C) Binding kinetics of UVR8^ValPro/AlaAla^ pre-monomerized by UV-B versus the COP1^Trp467Ala^ WD40 domain obtained by GCI experiments. Sensorgrams of UVR8 injected are shown in red, with their respective 1:1 binding model fits in black. The following amounts were typically used: ligand – COP1 (2000 pg/mm^2^); analyte – UVR8^VaPro/AlaAla^ +UV-B (highest concentration 2 μM). *k_a_* = association rate constant, *k_d_* = dissociation rate constant, *K_d_* = dissociation constant.

**Figure S10:**
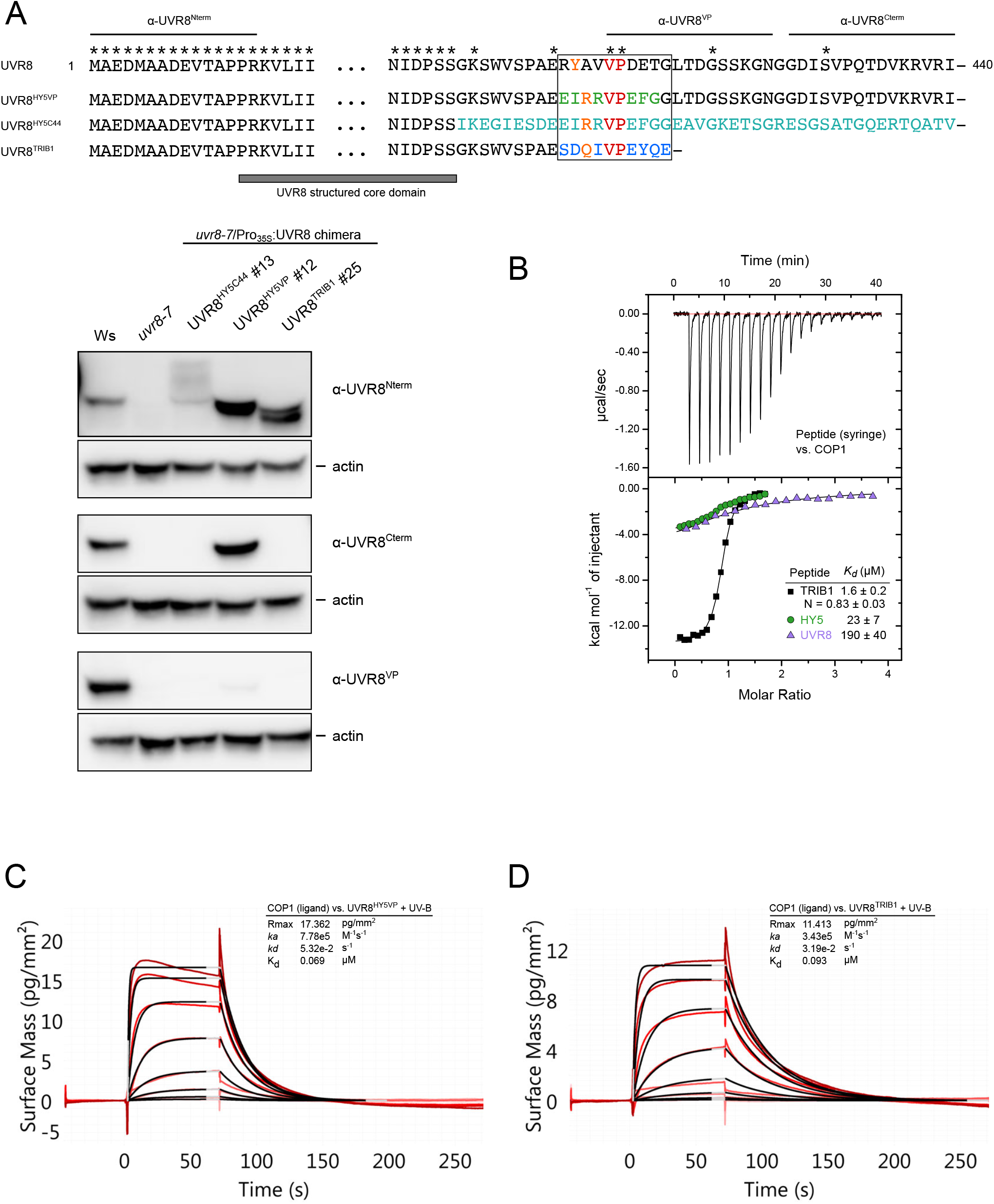
UVR8 chimeras can bind the COP1 WD40 domain. (A) Immunoblot analysis of UVR8 and actin (loading control) protein levels in 7-day-old wild-type (Ws), *uvr8-7* and *uvr8-7/Pro_35S_:UVR8^HY5C44^, uvr8-7/Pro_35S_:UVR8^HY5VP^* and *uvr8-7/Pro_35S_: UVR8_Trib_* seedlings. Specific antibodies against UVR8^1-15^ (α-UVR8^Nterm^), UVR8^410-424^ (α-UVR8^VP^) and UVR8^426-440^ (α-UVR8^Cterm^) were used. (B) ITC experiment between the TRIB1 VP-peptide versus the COP1 WD40 domain. Integrated heats are shown in solid, black squares. For comparison, ITC experiments between the UVR8 and HY5 VP peptides (from Figure 1B) versus the COP1 WD40 domain are shown in purple triangles and green circles, respectively. The following concentrations were typically used (titrant into cell): TRIB1 – COP1 (1000 μM in 175 μM). The inset shows the dissociation constant (*K_d_*), stoichiometry of binding (N) (± standard deviation). (C,D) Binding kinetics of the (C) UVR8^HY5C44^ or (D) UVR8^TRIB1VP^ chimeras pre-monomerized by UV-B versus COP1 obtained by GCI experiments. Sensorgrams of UVR8^HY5C44^ injected are shown in red, with their respective 1:1 binding model fits in black. The following amounts were typically used: ligand – COP1 (2000 pg/mm^2^); analyte – UVR8 chimeras (highest concentration 2 μM). *k_a_* = association rate constant, *k_d_* = dissociation rate constant, *K_d_* = dissociation constant.

**Figure S11:**
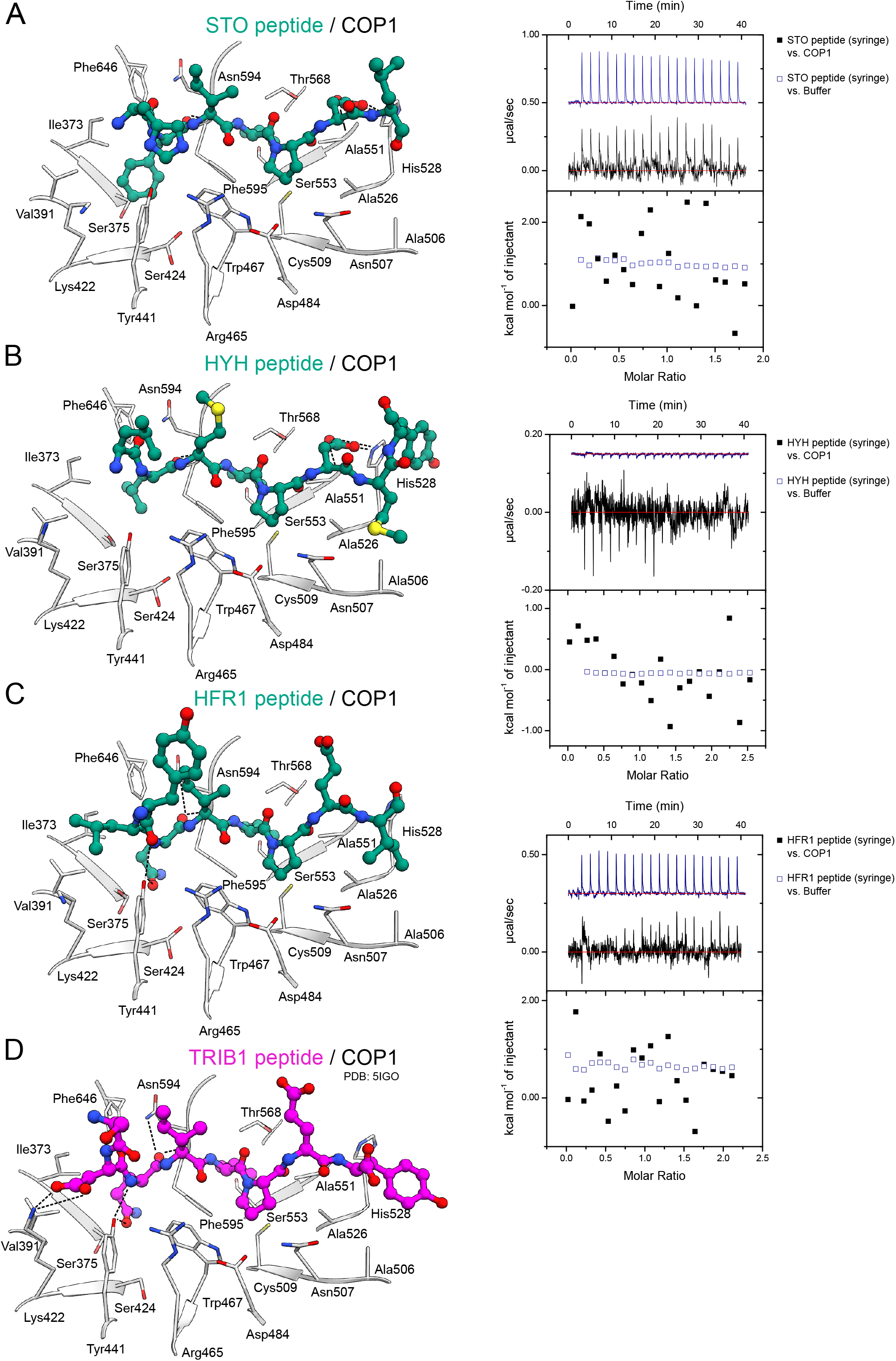
Various VP-peptides bind the COP1 WD40 domain. (A-D) (left) Crystal structure of the indicated peptide bound to the COP1 WD40 domain. The peptide is depicted in ball-and-stick representation. Selected residues from the COP1 WD40 domain are depicted in gray in stick representation. The TRIB1 – COP1 structure is from PDB-ID: 5IGO. (right) ITC assays between the indicated peptide versus the COP1 WD40 domain or buffer. The following concentrations were typically used (titrant into cell): STO – COP1 (1500 μM in 150 μM); HYH – COP1 (1500 μM in 125 μM); HFR1 – COP1 (1250 μM in 125 μM). The inset shows the dissociation constant (*K_d_*), stoichiometry of binding (N). (± standard deviation) See Figure S10B for the ITC experiment between the TRIB1 peptide versus the COP1 WD40 domain.

**Figure S12:**
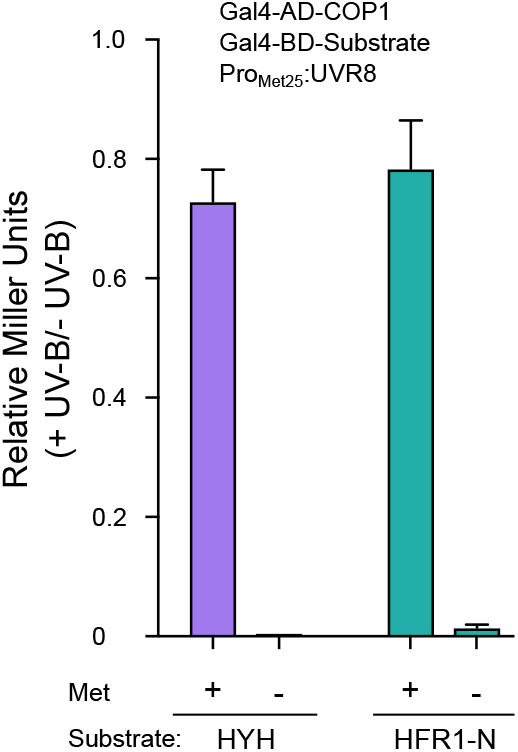
UVR8 can compete for COP1 binding against other COP1 interactors. Yeast 3-hybrid analysis of the COP1 - HYH, COP1 - HFR1 and COP1 - CRY1 interactions in the presence of UVR8. Normalized Miller Units were calculated as a ratio of β-galactosidase activity in yeast grown under UV-B versus yeast grown without UV-B. Additionally, normalized Miller Units here are reported separately for yeast grown on media without or with 1 mM methionine, corresponding to induction (− Met) or repression (+ Met) of *Met25* promoter-driven UVR8 expression, respectively. Means and SEM for 3 biological repetitions are shown. AD, Activation domain; BD, DNA binding domain; Met, methionine.

**Figure S13:**
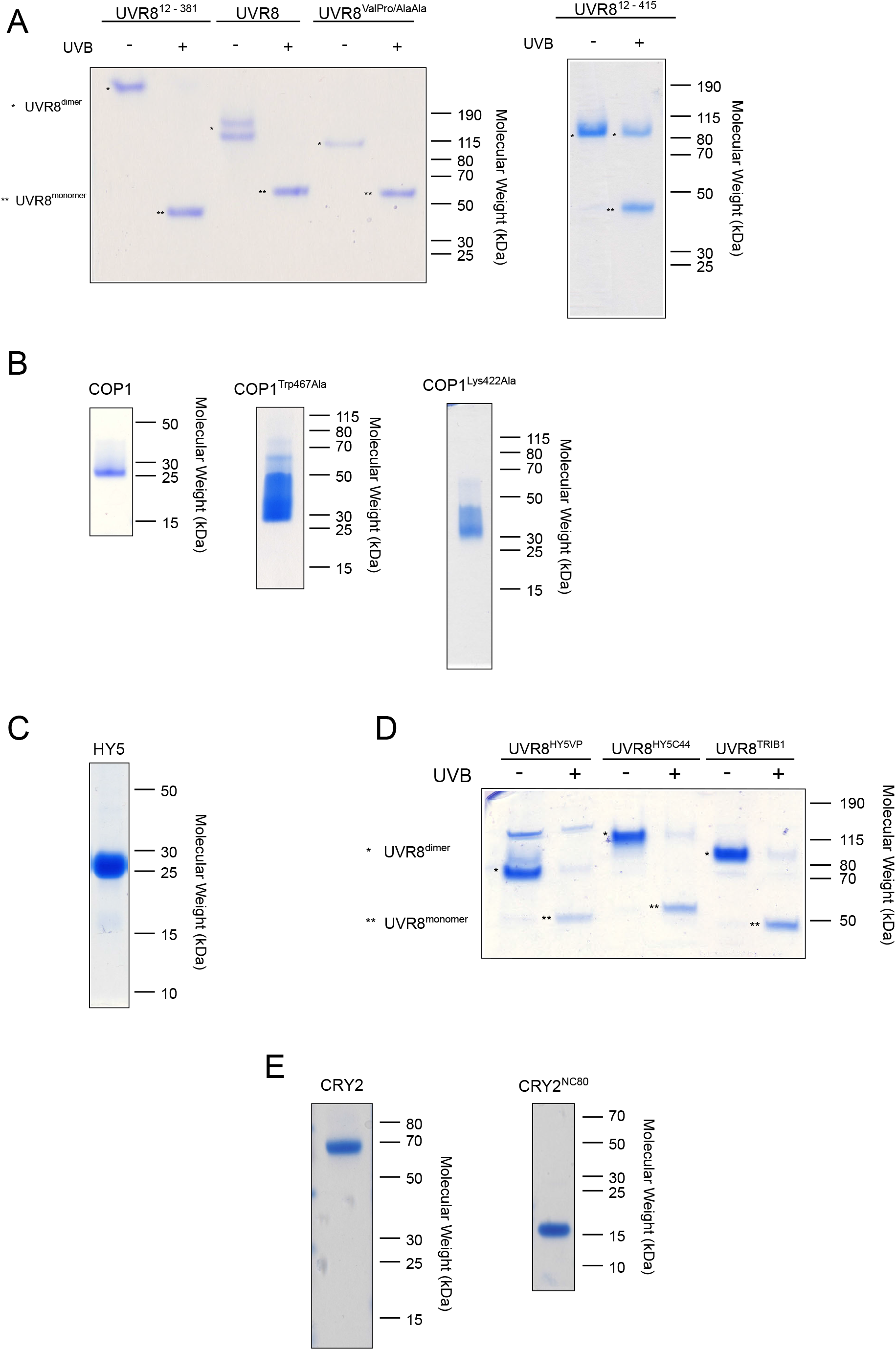
Coomassie-stained 10% SDS-PAGE gels of purified proteins show high purity. (A) Proteins used in Figure 2 and S6–9. (B) Proteins used in Figures 1–5, S1, S3, S6–11. (C) Proteins used in Figure 2. (D) Proteins used in Figure 3 and S10. (E) Proteins used in Figure 5.

## References

Adams, P.D., Afonine, P.V., Bunkóczi, G., Chen, V.B., Davis, I.W., Echols, N., Headd, J.J., Hung, L.-W., Kapral, G.J., Grosse-Kunstleve, R.W., et al. (2010). PHENIX: a comprehensive Python-based system for macromolecular structure solution. Acta Crystallogr. D Biol. Crystallogr. 66, 213–221.

Ahmad, M., and Cashmore, A.R. (1993). HY4 gene of A. thaliana encodes a protein with characteristics of a blue-light photoreceptor. Nature 366, 162–166.

Ahmad, M., Lin, C., and Cashmore, A.R. (1995). Mutations throughout an Arabidopsis blue-light photoreceptor impair blue-light-responsive anthocyanin accumulation and inhibition of hypocotyl elongation. Plant J. 8, 653–658.

Ang, L.-H., Chattopadhyay, S., Wei, N., Oyama, T., Okada, K., Batschauer, A., and Deng, X.-W. (1998). Molecular Interaction between COP1 and HY5 Defines a Regulatory Switch for Light Control of Arabidopsis Development. Mol Cell 1, 213–222.

von Arnim, A.G., and Deng, X.-W. (1994). Light inactivation of arabidopsis photomorphogenic repressor COP1 involves a cell-specific regulation of its nucleocytoplasmic partitioning. Cell 79, 1035–1045.

Arongaus, A.B., Chen, S., Pireyre, M., Glöckner, N., Galvão, V.C., Albert, A., Winkler, J.B., Fankhauser, C., Harter, K., and Ulm, R. (2018). Arabidopsis RUP2 represses UVR8-mediated flowering in noninductive photoperiods. Genes Dev. 32, 1332–1343.

Binkert, M., Kozma-Bognár, L., Terecskei, K., De Veylder, L., Nagy, F., and Ulm, R. (2014). UV-B-responsive association of the Arabidopsis bZIP transcription factor ELONGATED HYPOCOTYL5 with target genes, including its own promoter. Plant Cell 26, 4200–4213.

Binkert, M., Crocco, C.D., Ekundayo, B., Lau, K., Raffelberg, S., Tilbrook, K., Yin, R., Chappuis, R., Schalch, T., and Ulm, R. (2016). Revisiting chromatin binding of the Arabidopsis UV-B photoreceptor UVR8. BMC Plant Biol. 16, 42.

Brown, B.A., Cloix, C., Jiang, G.H., Kaiserli, E., Herzyk, P., Kliebenstein, D.J., and Jenkins, G.I. (2005). A UV-B-specific signaling component orchestrates plant UV protection. PNAS 102, 18225–18230.

Camacho, I.S., Theisen, A., Johannissen, L.O., Díaz-Ramos, L.A., Christie, J.M., Jenkins, G.I., Bellina, B., Barran, P., and Jones, A.R. (2019). Native mass spectrometry reveals the conformational diversity of the UVR8 photoreceptor. PNAS 116, 1116–1125.

Chen, H., Huang, X., Gusmaroli, G., Terzaghi, W., Lau, O.S., Yanagawa, Y., Zhang, Y., Li, J., Lee, J.-H., Zhu, D., et al. (2010). Arabidopsis CULLIN4-Damaged DNA Binding Protein 1 Interacts with CONSTITUTIVELY PHOTOMORPHOGENIC1-SUPPRESSOR OF PHYA Complexes to Regulate Photomorphogenesis and Flowering Time. Plant Cell 22, 108–123.

Christie, J.M., Arvai, A.S., Baxter, K.J., Heilmann, M., Pratt, A.J., O’Hara, A., Kelly, S.M., Hothorn, M., Smith, B.O., Hitomi, K., et al. (2012). Plant UVR8 Photoreceptor Senses UV-B by Tryptophan-Mediated Disruption of Cross-Dimer Salt Bridges. Science 335, 1492–1496.

Cloix, C., Kaiserli, E., Heilmann, M., Baxter, K.J., Brown, B.A., O’Hara, A., Smith, B.O., Christie, J.M., and Jenkins, G.I. (2012). C-terminal region of the UV-B photoreceptor UVR8 initiates signaling through interaction with the COP1 protein. PNAS 109, 16366–16370.

Clough, S.J., and Bent, A.F. (1998). Floral dip: a simplified method for Agrobacterium-mediated transformation of Arabidopsis thaliana. Plant J. 16, 735–743.

Datta, S., Hettiarachchi, G.H.C.M., Deng, X.-W., and Holm, M. (2006). Arabidopsis CONSTANS-LIKE3 Is a Positive Regulator of Red Light Signaling and Root Growth. Plant Cell 18, 70–84.

Deng, X.W., Caspar, T., and Quail, P.H. (1991). cop1: a regulatory locus involved in light-controlled development and gene expression in Arabidopsis. Genes Dev. 5, 1172–1182.

Deng, X.-W., Matsui, M., Wei, N., Wagner, D., Chu, A.M., Feldmann, K.A., and Quail, P.H. (1992). COP1, an arabidopsis regulatory gene, encodes a protein with both a zinc-binding motif and a Gβ homologous domain. Cell 71, 791–801.

Duek, P.D., Elmer, M.V., van Oosten, V.R., and Fankhauser, C. (2004). The degradation of HFR1, a putative bHLH class transcription factor involved in light signaling, is regulated by phosphorylation and requires COP1. Curr. Biol. 14, 2296–2301.

Durzynska, I., Xu, X., Adelmant, G., Ficarro, S.B., Marto, J.A., Sliz, P., Uljon, S., and Blacklow, S.C. (2017). STK40 Is a Pseudokinase that Binds the E3 Ubiquitin Ligase COP1. Structure 25, 287–294.

Emsley, P., and Cowtan, K. (2004). Coot: model-building tools for molecular graphics. Acta Cryst D 60, 2126–2132.

Favory, J.-J., Stec, A., Gruber, H., Rizzini, L., Oravecz, A., Funk, M., Albert, A., Cloix, C., Jenkins, G.I., Oakeley, E.J., et al. (2009). Interaction of COP1 and UVR8 regulates UV-B-induced photomorphogenesis and stress acclimation in Arabidopsis. EMBO J. 28, 591–601.

Gibson, D.G., Young, L., Chuang, R.-Y., Venter, J.C., Hutchison, C.A., and Smith, H.O. (2009). Enzymatic assembly of DNA molecules up to several hundred kilobases. Nat Meth 6, 343–345.

Gietz, R.D. (2014). Yeast transformation by the LiAc/SS carrier DNA/PEG method. Methods Mol. Biol. 1205, 1–12.

Gommers, C.M.M., and Monte, E. (2018). Seedling Establishment: A Dimmer Switch-Regulated Process between Dark and Light Signaling. Plant Physiol. 176, 1061–1074.

Gruber, H., Heijde, M., Heller, W., Albert, A., Seidlitz, H.K., and Ulm, R. (2010). Negative feedback regulation of UV-B–induced photomorphogenesis and stress acclimation in Arabidopsis. PNAS 107, 20132–20137.

Guo, H., Yang, H., Mockler, T.C., and Lin, C. (1998). Regulation of flowering time by Arabidopsis photoreceptors. Science 279, 1360–1363.

Hardtke, C.S., Gohda, K., Osterlund, M.T., Oyama, T., Okada, K., and Deng, X.W. (2000). HY5 stability and activity in Arabidopsis is regulated by phosphorylation in its COP1 binding domain. EMBO J. 19, 4997–5006.

Harper, J.W., Adami, G.R., Wei, N., Keyomarsi, K., and Elledge, S.J. (1993). The p21 Cdk-interacting protein Cip1 is a potent inhibitor of G1 cyclin-dependent kinases. Cell 75, 805–816.

Heijde, M., and Ulm, R. (2013). Reversion of the Arabidopsis UV-B photoreceptor UVR8 to the homodimeric ground state. PNAS 110, 1113–1118.

Heijde, M., Binkert, M., Yin, R., Ares-Orpel, F., Rizzini, L., Van De Slijke, E., Persiau, G., Nolf, J., Gevaert, K., De Jaeger, G., et al. (2013). Constitutively active UVR8 photoreceptor variant in Arabidopsis. PNAS 110, 20326–20331.

Heilmann, M., Velanis, C.N., Cloix, C., Smith, B.O., Christie, J.M., and Jenkins, G.I. (2016). Dimer/monomer status and in vivo function of salt-bridge mutants of the plant UV-B photoreceptor UVR8. Plant J. 88, 71–81.

Hoecker, U. (2017). The activities of the E3 ubiquitin ligase COP1/SPA, a key repressor in light signaling. Curr. Opin. Plant Biol. 37, 63–69.

Hoecker, U., and Quail, P.H. (2001). The Phytochrome A-specific Signaling Intermediate SPA1 Interacts Directly with COP1, a Constitutive Repressor of Light Signaling inArabidopsis. JBC 276, 38173–38178.

Holm, M., Hardtke, C.S., Gaudet, R., and Deng, X.-W. (2001). Identification of a structural motif that confers specific interaction with the WD40 repeat domain of Arabidopsis COP1. EMBO J. 20, 118–127.

Holm, M., Ma, L.-G., Qu, L.-J., and Deng, X.-W. (2002). Two interacting bZIP proteins are direct targets of COP1-mediated control of light-dependent gene expression in Arabidopsis. Genes Dev. 16, 1247–1259.

Holtkotte, X., Ponnu, J., Ahmad, M., and Hoecker, U. (2017). The blue light-induced interaction of cryptochrome 1 with COP1 requires SPA proteins during Arabidopsis light signaling. PLOS Genetics 13, e1007044.

Huang, X., Ouyang, X., Yang, P., Lau, O.S., Chen, L., Wei, N., and Deng, X.W. (2013). Conversion from CUL4-based COP1–SPA E3 apparatus to UVR8–COP1–SPA complexes underlies a distinct biochemical function of COP1 under UV-B. PNAS 110, 16669–16674.

Jang, I.-C., Yang, J.-Y., Seo, H.S., and Chua, N.-H. (2005). HFR1 is targeted by COP1 E3 ligase for post-translational proteolysis during phytochrome A signaling. Genes Dev. 19, 593–602.

Jang, I.-C., Henriques, R., Seo, H.S., Nagatani, A., and Chua, N.-H. (2010). Arabidopsis PHYTOCHROME INTERACTING FACTOR Proteins Promote Phytochrome B Polyubiquitination by COP1 E3 Ligase in the Nucleus. Plant Cell 22, 2370–2383.

Jang, S., Marchal, V., Panigrahi, K.C.S., Wenkel, S., Soppe, W., Deng, X.-W., Valverde, F., and Coupland, G. (2008). Arabidopsis COP1 shapes the temporal pattern of CO accumulation conferring a photoperiodic flowering response. EMBO J. 27, 1277–1288.

Jenkins, G.I. (2017). Photomorphogenic responses to ultraviolet-B light. Plant Cell Environ. 40, 2544–2557.

Kabsch, W. (1993). Automatic processing of rotation diffraction data from crystals of initially unknown symmetry and cell constants. J. Appl. Crystallogr. 26, 795–800.

Karimi, M., Inzé, D., and Depicker, A. (2002). GATEWAY vectors for Agrobacterium-mediated plant transformation. Trends Plant Sci. 7, 193–195.

Khanna, R., Kronmiller, B., Maszle, D.R., Coupland, G., Holm, M., Mizuno, T., and Wu, S.-H. (2009). The Arabidopsis B-Box Zinc Finger Family. Plant Cell 21, 3416–3420.

Kliebenstein, D.J., Lim, J.E., Landry, L.G., and Last, R.L. (2002). Arabidopsis UVR8 Regulates Ultraviolet-B Signal Transduction and Tolerance and Contains Sequence Similarity to Human Regulator of Chromatin Condensation 1. Plant Physiol. 130, 234–243.

Kung, J.E., and Jura, N. (2019). The pseudokinase TRIB1 toggles an intramolecular switch to regulate COP1 nuclear export. EMBO J. 38.

Lau, O.S., and Deng, X.W. (2012). The photomorphogenic repressors COP1 and DET1: 20 years later. Trends Plant Sci. 17, 584–593.

Lian, H.-L., He, S.-B., Zhang, Y.-C., Zhu, D.-M., Zhang, J.-Y., Jia, K.-P., Sun, S.-X., Li, L., and Yang, H.-Q. (2011). Blue-light-dependent interaction of cryptochrome 1 with SPA1 defines a dynamic signaling mechanism. Genes Dev. 25, 1023–1028.

Lin, C., and Shalitin, D. (2003). Cryptochrome Structure and Signal Transduction. Annu. Rev. Plant Biol. 54, 469–496.

Lin, F., Jiang, Y., Li, J., Yan, T., Fan, L., Liang, J., Chen, Z.J., Xu, D., and Deng, X.W. (2018). B-BOX DOMAIN PROTEIN28 Negatively Regulates Photomorphogenesis by Repressing the Activity of Transcription Factor HY5 and Undergoes COP1-Mediated Degradation. Plant Cell 30, 2006–2019.

Liu, H., and Naismith, J.H. (2008). An efficient one-step site-directed deletion, insertion, single and multiple-site plasmid mutagenesis protocol. BMC Biotechnology 8, 91.

Liu, B., Zuo, Z., Liu, H., Liu, X., and Lin, C. (2011). Arabidopsis cryptochrome 1 interacts with SPA1 to suppress COP1 activity in response to blue light. Genes Dev. 25, 1029–1034.

Liu, L.-J., Zhang, Y.-C., Li, Q.-H., Sang, Y., Mao, J., Lian, H.-L., Wang, L., and Yang, H.-Q. (2008). COP1-Mediated Ubiquitination of CONSTANS Is Implicated in Cryptochrome Regulation of Flowering in Arabidopsis. Plant Cell 20, 292–306.

Livak, K.J., and Schmittgen, T.D. (2001). Analysis of relative gene expression data using real-time quantitative PCR and the 2(-Delta Delta C(T)) Method. Methods 25, 402–408.

Lu, X.-D., Zhou, C.-M., Xu, P.-B., Luo, Q., Lian, H.-L., and Yang, H.-Q. (2015). Red-light-dependent interaction of phyB with SPA1 promotes COP1-SPA1 dissociation and photomorphogenic development in Arabidopsis. Mol Plant 8, 467–478.

Marrocco, K., Zhou, Y., Bury, E., Dieterle, M., Funk, M., Genschik, P., Krenz, M., Stolpe, T., and Kretsch, T. (2006). Functional analysis of EID1, an F-box protein involved in phytochrome A-dependent light signal transduction. Plant J. 45, 423–438.

McCoy, A.J., Grosse-Kunstleve, R.W., Adams, P.D., Winn, M.D., Storoni, L.C., and Read, R.J. (2007). Phaser crystallographic software. J. Appl. Crystallogr. 40, 658–674.

McNellis, T.W., Arnim, A.G. von, Araki, T., Komeda, Y., Miséra, S., and Deng, X.W. (1994). Genetic and molecular analysis of an allelic series of cop1 mutants suggests functional roles for the multiple protein domains. Plant Cell 6, 487–500.

Meijering, E., Jacob, M., Sarria, J.-C.F., Steiner, P., Hirling, H., and Unser, M. (2004). Design and validation of a tool for neurite tracing and analysis in fluorescence microscopy images. Cytometry A 58, 167–176.

Möller-Steinbach, Y., Alexandre, C., and Hennig, L. (2010). Flowering time control. Methods Mol. Biol. 655, 229–237.

Müller, P., and Bouly, J.-P. (2015). Searching for the mechanism of signalling by plant photoreceptor cryptochrome. FEBS Letters 589, 189–192.

Murshudov, G.N., Skubák, P., Lebedev, A.A., Pannu, N.S., Steiner, R.A., Nicholls, R.A., Winn, M.D., Long, F., and Vagin, A.A. (2011). REFMAC5 for the refinement of macromolecular crystal structures. Acta Cryst D 67, 355–367.

Newton, K., Dugger, D.L., Sengupta-Ghosh, A., Ferrando, R.E., Chu, F., Tao, J., Lam, W., Haller, S., Chan, S., Sa, S., et al. (2018). Ubiquitin ligase COP1 coordinates transcriptional programs that control cell type specification in the developing mouse brain. PNAS 115, 11244–11249.

Oravecz, A., Baumann, A., Máté, Z., Brzezinska, A., Molinier, J., Oakeley, E.J., Adám, E., Schäfer, E., Nagy, F., and Ulm, R. (2006). CONSTITUTIVELY PHOTOMORPHOGENIC1 is required for the UV-B response in Arabidopsis. Plant Cell 18, 1975–1990.

Ordoñez-Herrera, N., Fackendahl, P., Yu, X., Schaefer, S., Koncz, C., and Hoecker, U. (2015). A cop1 spa mutant deficient in COP1 and SPA proteins reveals partial co-action of COP1 and SPA during Arabidopsis post-embryonic development and photomorphogenesis. Mol Plant 8, 479–481.

Ordoñez-Herrera, N., Trimborn, L., Menje, M., Henschel, M., Robers, L., Kaufholdt, D., Hänsch, R., Adrian, J., Ponnu, J., and Hoecker, U. (2018). The Transcription Factor COL12 Is a Substrate of the COP1/SPA E3 Ligase and Regulates Flowering Time and Plant Architecture. Plant Physiol. 176, 1327–1340.

Osterlund, M.T., Hardtke, C.S., Wei, N., and Deng, X.W. (2000). Targeted destabilization of HY5 during light-regulated development of Arabidopsis. Nature 405, 462–466.

Oyama, T., Shimura, Y., and Okada, K. (1997). The Arabidopsis HY5 gene encodes a bZIP protein that regulates stimulus-induced development of root and hypocotyl. Genes Dev. 11, 2983–2995.

Pettersen, E.F., Goddard, T.D., Huang, C.C., Couch, G.S., Greenblatt, D.M., Meng, E.C., and Ferrin, T.E. (2004). UCSF Chimera-a visualization system for exploratory research and analysis. J Comput Chem 25, 1605–1612.

Pham, V.N., Xu, X., and Huq, E. (2018). Molecular bases for the constitutive photomorphogenic phenotypes in Arabidopsis. Development 145, dev169870.

Podolec, R., and Ulm, R. (2018). Photoreceptor-mediated regulation of the COP1/SPA E3 ubiquitin ligase. Current Opinion in Plant Biology 45, 18–25.

Putterill, J., Robson, F., Lee, K., Simon, R., and Coupland, G. (1995). The CONSTANS gene of arabidopsis promotes flowering and encodes a protein showing similarities to zinc finger transcription factors. Cell 80, 847–857.

Ren, H., Han, J., Yang, P., Mao, W., Liu, X., Qiu, L., Qian, C., Liu, Y., Chen, Z., Ouyang, X., et al. (2019). Two E3 ligases antagonistically regulate the UV-B response in Arabidopsis. PNAS 201816268.

Rizzini, L., Favory, J.-J., Cloix, C., Faggionato, D., O’Hara, A., Kaiserli, E., Baumeister, R., Schäfer, E., Nagy, F., Jenkins, G.I., et al. (2011). Perception of UV-B by the Arabidopsis UVR8 Protein. Science 332, 103–106.

Saijo, Y., Sullivan, J.A., Wang, H., Yang, J., Shen, Y., Rubio, V., Ma, L., Hoecker, U., and Deng, X.W. (2003). The COP1-SPA1 interaction defines a critical step in phytochrome A-mediated regulation of HY5 activity. Genes Dev. 17, 2642–2647.

Seo, H.S., Yang, J.-Y., Ishikawa, M., Bolle, C., Ballesteros, M.L., and Chua, N.-H. (2003). LAF1 ubiquitination by COP1 controls photomorphogenesis and is stimulated by SPA1. Nature 423, 995–999.

Sheerin, D.J., Menon, C., Oven-Krockhaus, S. zur, Enderle, B., Zhu, L., Johnen, P., Schleifenbaum, F., Stierhof, Y.-D., Huq, E., and Hiltbrunner, A. (2015). Light-Activated Phytochrome A and B Interact with Members of the SPA Family to Promote Photomorphogenesis in Arabidopsis by Reorganizing the COP1/SPA Complex. Plant Cell 27, 189–201.

Stelzl, U., Worm, U., Lalowski, M., Haenig, C., Brembeck, F.H., Goehler, H., Stroedicke, M., Zenkner, M., Schoenherr, A., Koeppen, S., et al. (2005). A human protein-protein interaction network: a resource for annotating the proteome. Cell 122, 957–968.

Uljon, S., Xu, X., Durzynska, I., Stein, S., Adelmant, G., Marto, J.A., Pear, W.S., and Blacklow, S.C. (2016). Structural Basis for Substrate Selectivity of the E3 Ligase COP1. Structure 24, 687–696.

Ulm, R., Baumann, A., Oravecz, A., Máté, Z., Ádám, É., Oakeley, E.J., Schäfer, E., and Nagy, F. (2004). Genome-wide analysis of gene expression reveals function of the bZIP transcription factor HY5 in the UV-B response of Arabidopsis. PNAS 101, 1397–1402.

Vandenbussche, F., Tilbrook, K., Fierro, A.C., Marchal, K., Poelman, D., Van Der Straeten, D., and Ulm, R. (2014). Photoreceptor-mediated bending towards UV-B in Arabidopsis. Mol Plant 7, 1041–1052.

Viczián, A., Ádám, É., Wolf, I., Bindics, J., Kircher, S., Heijde, M., Ulm, R., Schäfer, E., and Nagy, F. (2012). A short amino-terminal part of Arabidopsis phytochrome A induces constitutive photomorphogenic response. Mol Plant 5, 629–641.

Vojtek, A.B., and Hollenberg, S.M. (1995). Ras-Raf interaction: two-hybrid analysis. Meth. Enzymol. 255, 331–342.

Wang, H., Ma, L.-G., Li, J.-M., Zhao, H.-Y., and Deng, X.W. (2001). Direct Interaction of Arabidopsis Cryptochromes with COP1 in Light Control Development. Science 294, 154–158.

Wang, Q., Zuo, Z., Wang, X., Liu, Q., Gu, L., Oka, Y., and Lin, C. (2018). Beyond the photocycle-how cryptochromes regulate photoresponses in plants? Curr. Opin. Plant Biol. 45, 120–126.

Wang, Z.-P., Xing, H.-L., Dong, L., Zhang, H.-Y., Han, C.-Y., Wang, X.-C., and Chen, Q.-J. (2015). Egg cell-specific promoter-controlled CRISPR/Cas9 efficiently generates homozygous mutants for multiple target genes in Arabidopsis in a single generation. Genome Biol. 16, 144.

Wickham, H. (2009). ggplot2: Elegant Graphics for Data Analysis (New York: Springer-Verlag).

Winn, M.D., Ballard, C.C., Cowtan, K.D., Dodson, E.J., Emsley, P., Evans, P.R., Keegan, R.M., Krissinel, E.B., Leslie, A.G.W., McCoy, A., et al. (2011). Overview of the CCP4 suite and current developments. Acta Cryst D 67, 235–242.

Wu, D., Hu, Q., Yan, Z., Chen, W., Yan, C., Huang, X., Zhang, J., Yang, P., Deng, H., Wang, J., et al. (2012). Structural basis of ultraviolet-B perception by UVR8. Nature 484, 214–219.

Wu, M., Farkas, D., Eriksson, L.A., and Strid, Å. (2019). Proline 411 biases the conformation of the intrinsically disordered plant UVR8 photoreceptor C27 domain altering the functional properties of the peptide. Sci Rep 9, 818.

Xu, D., Jiang, Y., Li, J., Lin, F., Holm, M., and Deng, X.W. (2016). BBX21, an Arabidopsis B-box protein, directly activates HY5 and is targeted by COP1 for 26S proteasome-mediated degradation. PNAS 113, 7655–7660.

Yan, H., Marquardt, K., Indorf, M., Jutt, D., Kircher, S., Neuhaus, G., and Rodríguez-Franco, M. (2011). Nuclear localization and interaction with COP1 are required for STO/BBX24 function during photomorphogenesis. Plant Physiol. 156, 1772–1782.

Yang, J., and Wang, H. (2006). The central coiled-coil domain and carboxyl-terminal WD-repeat domain of Arabidopsis SPA1 are responsible for mediating repression of light signaling. Plant J. 47, 564–576.

Yang, H.-Q., Wu, Y.-J., Tang, R.-H., Liu, D., Liu, Y., and Cashmore, A.R. (2000). The C Termini of Arabidopsis Cryptochromes Mediate a Constitutive Light Response. Cell 103, 815–827.

Yang, H.-Q., Tang, R.-H., and Cashmore, A.R. (2001). The Signaling Mechanism of Arabidopsis CRY1 Involves Direct Interaction with COP1. Plant Cell 13, 2573–2587.

Yang, J., Lin, R., Sullivan, J., Hoecker, U., Liu, B., Xu, L., Deng, X.W., and Wang, H. (2005). Light Regulates COP1-Mediated Degradation of HFR1, a Transcription Factor Essential for Light Signaling in Arabidopsis. Plant Cell 17, 804–821.

Yang, L., Mo, W., Yu, X., Yao, N., Zhou, Z., Fan, X., Zhang, L., Piao, M., Li, S., Yang, D., et al. (2018). Reconstituting Arabidopsis CRY2 Signaling Pathway in Mammalian Cells Reveals Regulation of Transcription by Direct Binding of CRY2 to DNA. Cell Reports 24, 585–593.e4.

Yin, R., and Ulm, R. (2017). How plants cope with UV-B: from perception to response. Curr. Opin. Plant Biol. 37, 42–48.

Yin, R., Messner, B., Faus-Kessler, T., Hoffmann, T., Schwab, W., Hajirezaei, M.-R., von Saint Paul, V., Heller, W., and Schäffner, A.R. (2012). Feedback inhibition of the general phenylpropanoid and flavonol biosynthetic pathways upon a compromised flavonol-3-O-glycosylation. J. Exp. Bot. 63, 2465–2478.

Yin, R., Arongaus, A.B., Binkert, M., and Ulm, R. (2015). Two distinct domains of the UVR8 photoreceptor interact with COP1 to initiate UV-B signaling in Arabidopsis. Plant Cell 27, 202–213.

Yu, X., Shalitin, D., Liu, X., Maymon, M., Klejnot, J., Yang, H., Lopez, J., Zhao, X., Bendehakkalu, K.T., and Lin, C. (2007). Derepression of the NC80 motif is critical for the photoactivation of Arabidopsis CRY2. PNAS 104, 7289–7294.

Yuan, T.-T., Xu, H.-H., Zhang, Q., Zhang, L.-Y., and Lu, Y.-T. (2018). The COP1 Target SHI-RELATED SEQUENCE5 Directly Activates Photomorphogenesis-Promoting Genes. Plant Cell 30, 2368–2382.

Zhu, D., Maier, A., Lee, J.-H., Laubinger, S., Saijo, Y., Wang, H., Qu, L.-J., Hoecker, U., and Deng, X.W. (2008). Biochemical Characterization of Arabidopsis Complexes Containing CONSTITUTIVELY PHOTOMORPHOGENIC1 and SUPPRESSOR OF PHYA Proteins in Light Control of Plant Development. Plant Cell 20, 2307–2323.

Zuo, Z., Liu, H., Liu, B., Liu, X., and Lin, C. (2011). Blue light-dependent interaction of CRY2 with SPA1 regulates COP1 activity and floral initiation in Arabidopsis. Curr. Biol. 21, 841–847.

